# A survey of dietary effects on tRNA abundance and modifications

**DOI:** 10.1101/2025.06.10.658430

**Authors:** Simeiyun Liu, Andrew D. Holmes, Ruoxia Zhao, Cassandra Herbert, Patrick A. Limbach, Todd M. Lowe, Upasna Sharma

**Affiliations:** Department of Molecular, Cell and Developmental Biology, University of California, Santa Cruz, California, 95064, United States; Department of Chemistry, University of Cincinnati, P.O. Box 210172, Cincinnati, Ohio 45221-0172, United States; Department of Biomedical Engineering, University of California, Santa Cruz, California, 95064, United States

## Abstract

Transfer RNAs (tRNAs) play a central role in protein translation and are increasingly recognized as dynamic regulators of gene expression. Both physiological and environmental signals can modulate tRNA abundance and chemical modifications, yet the impact of dietary cues on the tRNA landscape remains poorly understood. Here, we investigated the effects of two distinct dietary interventions—low-protein and high-fat diets—on tRNA abundance and modification profiles across multiple mouse tissues. We conducted a comprehensive analysis of tRNA abundance and modification changes in response to these nutritional challenges using RNA mass spectrometry and Ordered Two-Template Relay sequencing (OTTR-seq), a modified-base-sensitive tRNA sequencing method. Our results reveal both shared and tissue-specific alterations in abundance and modifications of specific nuclear and mitochondrial genome-encoded tRNAs in response to dietary conditions at isotype, isoacceptor, and isodecoder levels. As many of the tissue-specific or diet-responsive tRNA modifications have been previously reported to affect decoding efficiency or translational fidelity, these results have implications for understanding translational adaptation in response to dietary conditions.

## Introduction

Transfer RNAs (tRNAs) play a central role in protein synthesis by decoding messenger RNA (mRNA) information into corresponding amino acid sequences, thereby translating the genetic code into functional proteins. Notably, most tRNAs are encoded by multiple genes, expanding the repertoire of tRNA molecules available for each type and allowing for fine-tuned regulation of translational dynamics. tRNAs are broadly categorized into isotypes (each charged with one of the 20 standard amino acids), isoacceptors (tRNAs with different anticodons that carry the same amino acid), and isodecoders (tRNAs sharing the same anticodon but differing in sequence elsewhere in the molecule). tRNAs are also among the most extensively modified RNA species in the cell, with over 100 distinct chemical modifications identified to date and an average of ∼13 modifications per tRNA molecule [1]. These post-transcriptional modifications are essential for proper tRNA folding, structural stability, and accurate decoding during translation.

Recent studies report that tRNA abundance varies across different mouse tissues [2–5]. Furthermore, many tRNA-modifying enzymes are expressed in a tissue-specific manner. Comparative analysis of the expression of RNA-modifying proteins across various tissues revealed that certain proteins are specifically expressed in the reproductive tract [6], including the testis and the epididymis− an organ where sperm undergo post-testicular maturation and become motile. For example, NSUN7, a 5-methylcytosine (m^5^C) modifying enzyme, is expressed during the late stages of spermatogenesis in the spermatocytes and spermatids [6] and is required for male fertility [7]. Another member of the family, NSUN2, is also required for the meiotic progression of germ cells in the testis [8]. NSUN2 and NSUN7 are enriched in the proximal region of the epididymis (caput epididymis) but not in the distal epididymis (cauda epididymis), indicating segment-specific expression of these proteins in the epididymis. Moreover, the epididymis was reported to be specifically enriched in two additional tRNA modifying enzymes: TRDMT1 (also known as DNMT2), a m^5^C methyltransferase known to modify position 38 in specific tRNAs, and METTL1, a N7-methylguanosine (m^7^G) tRNA methyltransferase [6]. Whether reproductive tract tissues have differential tRNA modification profiles compared to other tissues remains unknown.

Stress and other external factors can also influence tRNA abundance and modifications [9–11]. For instance, high reactive oxygen species conditions in *E. coli* cells can lead to elevated tRNA Tyr-QUA-II m^5^C levels at position 49 [12]. The addition of hydrogen peroxide (H₂O₂) to yeast cells has been shown to increase the levels of specific posttranscriptional modifications, including m⁵C, N², N²-dimethylguanosine (m²,²G), and 2’-O-methylcytidine (Cm) [11]. Similarly, arsenite exposure increased queuosine levels, which were associated with the upregulation of proteins encoded by codon-biased genes involved in energy metabolism [13]. tRNA thiolation is downregulated in response to sulfur starvation [14]. Moreover, the gut microbiome has been reported to regulate tRNA abundance and modifications in the host in a tissue-specific manner [15]. Conversely, diet can influence tRNA abundance and modifications in the gut microbiome communities [16]. In addition to their role in translation, tRNAs can be cleaved to generate tRNA-derived RNAs (tDRs) [17], also known as tRNA-derived small RNAs (tsRNAs) or tRNA fragments (tRFs), which have been proposed to play key roles in various cellular processes [18, 19]. Exposure to low-protein or high-fat diets leads to alterations in the levels of specific tDRs in sperm [20, 21]. Whether diet alters the levels and modifications of mature tRNAs remains unknown. Determining the effects of various internal and external factors on tRNAs is important as alterations in tRNA abundance and/or modifications can lead to selective translation of codon-biased mRNAs [11].

Major advancements in identifying and understanding the dynamic nature of tRNA modifications were made possible through the development of liquid chromatography-tandem mass spectrometry (LC-MS/MS) methods to measure RNA nucleotide modifications accurately [22, 23]. Despite these advances, mass-spectrometry-based methods cannot provide single-molecule-level information. On the other hand, accurate measurement of over 400 distinct mouse tRNA molecules by deep sequencing has been challenging due to the extensive structural and chemical modifications in tRNAs and an inability to distinguish between isodecoders. Overcoming these challenges, various new methods for mature tRNA sequencing and analysis have been developed recently [24], including demethylase-thermostable group II intron RT tRNA sequencing (DM-tRNA-seq) [25], AlkB-facilitated RNA methylation sequencing (ARM-seq) [26], Hydro-tRNAseq [27], Y-shaped adapter-ligated mature tRNA sequencing (YAMAT-seq) [28], long hairpin oligonucleotide-based tRNA high-throughput sequencing (LOTTE tRNAseq) [29], quantitative mature tRNA sequencing (QuantM-tRNAseq) [2], modification-induced misincorporation tRNA sequencing (mim-tRNAseq) [30], Nano-tRNAseq [31], and ligation-independent detection of all types of RNA (LIDAR-seq) [32].

Some of these methods use enzymatic treatment to remove modifications that pause or stop reverse transcription (RT), including N^1^-methyladenosine (m1A), N*^3^*-methylcytidine (m^3^C), and N^1^-methylguanosine (m^1^G) [2, 25, 26, 33, 34]. In addition, high processivity RT enzymes have been used to allow sequencing of highly modified tRNAs. High processivity RT enzymes often misincorporate nucleotides at some modified bases, thus providing single-nucleotide resolution of putative modified positions in tRNAs. One such method, DM-tRNA-seq, involves treating tRNA with the *E. Coli* demethylase AlkB in combination with the TGIRT enzyme to allow more efficient full-length tRNA sequencing [25]. Recently, Ordered Two-Template Relay sequencing (OTTR-seq) was developed, which uses an engineered recombinant non-LTR retroelement RT that adds both adaptors to the cDNA in a single RT step by template jumping [35], allowing the generation of low-bias small RNA sequencing libraries. This method has been successfully used to sequence full-length tRNAs from yeast and mice [36]. Importantly, OTTR-seq provides site-specific information on modified nucleotides through reverse transcriptase misincorporation signatures, overcoming prior technical challenges due to extensive base modifications in tRNAs and the difficulty of adapter ligation [35, 36].

While it is increasingly appreciated that various internal and external factors alter tRNA abundance and modifications, there is limited understanding of how dietary factors influence cellular tRNAs. Here, we used a combination of RNA mass spectrometry and modification-sensitive small RNA sequencing (OTTR-seq) to examine changes in tRNA transcript abundance and modifications in somatic (liver and heart) and male reproductive (testis, epididymis, and sperm) tissues from mice subjected to high-fat or low-protein diets. Our findings highlight that tRNAs and their modifications respond to diet in a tissue-specific manner, with differential nuclear and mitochondrial tRNAs responses. These results have implications for understanding translational regulation and metabolic adaptation in response to dietary conditions.

## Results

### The liver and epididymis tissues display differential tRNA modification levels and respond uniquely to dietary changes

To examine if the levels of tRNA modifications are regulated in a tissue-specific manner, we examined the abundance of specific modifications in tRNAs and small RNAs (sRNA) isolated from the epididymis and liver tissues using LC-MS/MS (**Figure 1A**). We focused on these two tissues to study RNA modifications in reproductive and non-reproductive tract tissues, given that specific RNA-modifying enzymes are more abundant in the reproductive tissues [6]. sRNAs ranging from 16 to 40 nucleotides (nts) (primarily consisting of tRFs) and RNAs ranging from 70 to 90 nts (corresponding to mature tRNA size) were isolated from total RNA by size fractionation on a polyacrylamide gel (**Figure S1A and S1B**) and processed for RNA nucleotide modification analysis using LC-MS/MS for 24 known RNA modifications (**Figure S1C**). 9 out of 24 (36%) modifications were differentially enriched between the two tissues (significance cut-off used: p value <0.05, fold change >1.5, two-sample t-test). For example, in the sRNA fraction, the levels of m^2^,^2^G were higher in the liver, while m^3^U were higher in the epididymis (**Figure 1B**). The 70-90nts fraction was primarily composed of mature tRNAs as determined by DM-tRNA-seq[37] (**Figure S1D**). In tRNAs, galactosyl-queuosine (galQ) and 2’-O-methyladenosine (Am) were higher in the epididymis, and 5,2’-O-dimethyluridine (m^5^Um) was relatively more abundant in the liver (**Figure 1C**). The differences in modification levels between the two tissues could be due to differences in their tRNA composition (**Figure S1E**), as individual tRNAs can carry unique combinations of modifications. However, in the case of galQ modification, higher galQ levels in the epididymis compared to the liver were not correlated with tRNA-Tyr abundance (the only tRNA known to have this modification [38]) (**Figure S1F**),

**Figure 1.**
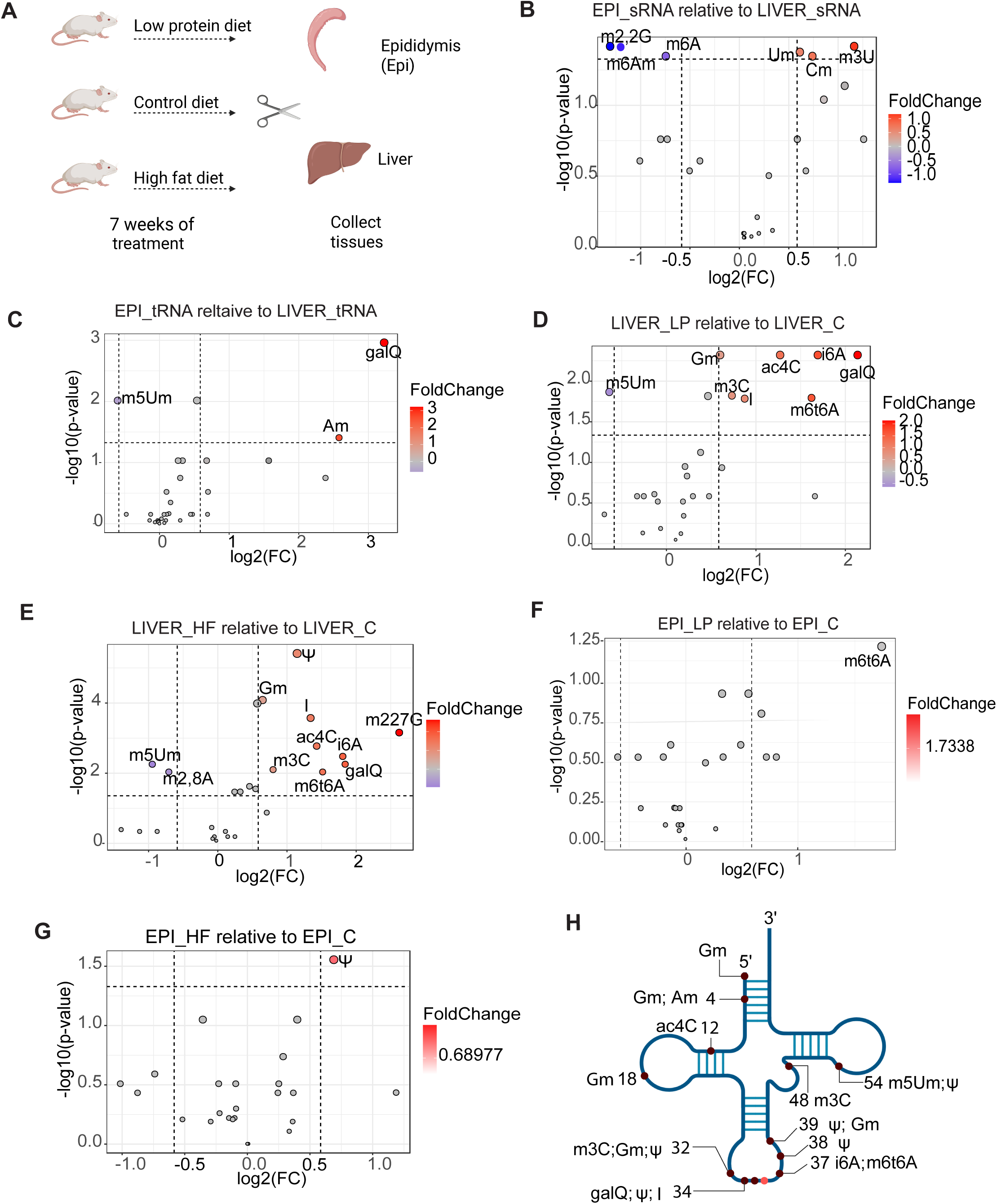
Tissue-specific and diet-induced alterations in tRNA modifications in the liver and epididymis. **A)** Schematic of the experimental design. 3-week-old male mice were fed one of three diets for 7 weeks: low-protein (LP), high-fat (HF), or control (C). Liver and epididymis (EPI) tissues were collected for downstream analyses. **B)** Volcano plot of small RNA (16 - 40nts) modifications in the epididymis relative to liver. Modifications detected at significantly differential levels between the two tissues are labeled on the plot (modifications that showed log2FC ≥ 0.585 and -log10(p-value) > 1.3). The color scale represents the fold change. Dot size reflects the -log10(p-value). n=6 replicates. **C)** Volcano plot showing significantly altered tRNA modifications (70 - 90nts) in the epididymis relative to the liver (n = 6). **D-G)** Volcano plots illustrating RNA modifications significantly affected by LP or HF diets in the liver and epididymis tissues (HF sample n = 3; control and LP sample n = 6). **H)** Schematic of tissue-specific and diet-responsive tRNA modifications identified by LC-MS/MS, annotated with positional information on the tRNA.

Next, we focused on the effects of diet on RNA modification levels in the epididymis and liver. We fed male mice either a low-protein diet (LP, 10% protein), a high-fat diet (HF, 60% fat), or a control diet (C, 18% protein, 7.1% fat) from weaning until 10 weeks of age **(Figure 1A)**. As expected, mice fed a high-fat diet gained significant weight compared to control diet-fed mice by the end of the intervention (**Figure S1G**). We did not observe any change in the weight of low-protein diet-fed mice compared to control animals, which is consistent with previous observations [20]. Modification levels in 16–40 nt sRNAs were unaffected by either LP or HF diets (data not shown). In contrast, in full-length tRNAs, 11 out of 24 modifications (46%) exhibited diet-induced alterations (significance cut-off used: p value <0.05, fold change >1.5, two-sample t-test; **Figure 1D–G**). The liver showed robust changes in response to dietary alterations, consistent with its central role in metabolism, digestion, and nutrient storage (**Figure 1D-E**). Interestingly, several RNA modifications were altered similarly under the two dietary conditions in the liver: galQ, N4-acetylcytidine (ac^4^C), N6-methyl-threonylcarbamoyl adenosine (m^6^t^6^A), m^3^C, 2’-O-methylguanosine (Gm), inosine (I), and N^6^-isopentenyladenosine (i^6^A) were upregulated, and m^5^Um was downregulated (**Figure 1D-E**). In addition to these shared changes with LP diet, HF livers also showed an increase in pseudouridine (Ψ) and N^2^, N^2^, N^7^-trimethylguanosine (m^2^,^2^,^7^G) and downregulation of 2,8-dimethyladenosine (m^2^,^8^A) (**Figure 1E**). These findings suggest that LP and HF diets impact overlapping and distinct pathways, leading to shared and unique alterations in hepatic tRNA modification landscapes. Additionally, the detection of alterations in m^2^,^8^A [39] and m^2^,^2^,^7^G [40] —which are typically enriched in rRNAs and snRNAs, respectively—suggests that dietary interventions may influence RNA modifications across multiple RNA species, not just tRNAs. The epididymis exhibited a modest response to dietary changes compared to the liver. In the epididymis, there was an increase in Ψ levels in the HF condition, with no change observed in response to LP diet (**Figure 1F-G**). As Ψ was upregulated in both the epididymis and liver tissues under HF dietary challenge, HF diet potentially drives systemic changes in some tRNA modifications.

These findings indicate that tRNA modifications are regulated in a tissue-specific manner, and dietary metabolites can drive systemic and tissue-specific changes in modification levels. The liver can modulate gene expression and enzyme activity in response to macronutrient composition and dietary stress to help cells adapt to metabolic demands [41]. Our data revealed that the liver also alters tRNA modification status in response to nutritional challenges, adding another level of gene regulation to help cells adapt to such stressors. As shown in **Figure 1H**, many RNA modifications altered by tissue type or dietary conditions are enriched around the anticodon region, a key site for translation regulation. These tRNA modification changes under dietary stress suggest that tRNAs may help tune translation in response to metabolic demands [42].

### tRNA transcript abundance is regulated in a tissue-specific manner

While LC-MS/MS remains the gold standard for quantifying RNA modifications with high sensitivity and precision, it does not provide information on the specific tRNA species that are differentially modified, limiting insight into the transcript-level resolution of these changes. Therefore, we next assessed tRNA abundance and modification status across multiple tissues at single-nucleotide resolution using OTTR-seq [36]. To more robustly capture diet-induced changes in tRNA abundance and modification, we further refined our experimental design for OTTR-seq (**Figure S2A**). *First*, we extended dietary exposure to six months to better reflect long-term nutritional effects. *Second*, we introduced a more appropriate control for the high-fat (HF) diet, ensuring that the HF diet and its corresponding control diet (C_HF) differed only in fat content. *Third*, rather than analyzing the epididymis as a whole (as done in LC-MS/MS analysis), we separately examined the proximal caput (CP) and distal cauda (CA) segments of the epididymis, which are known to exhibit distinct gene expression patterns—including differential expression of RNA-modifying enzymes—and play specialized roles in sperm maturation [6, 43–45]. *Fourth*, we broadened the scope of our tRNA expression and modification analysis to include additional tissues, namely the heart, testis, and mature sperm, to assess systemic and germline-specific responses to dietary interventions.

Mice were subjected to the dietary interventions described above, total RNA was isolated from tissues, sequencing libraries were generated using OTTR-seq, and data were analyzed using tRNA Analysis of eXpression (tRAX), an analytical pipeline designed for comprehensive profiling of tRNAs and modified nucleotides [46] (**Figure S2B-C**). The relative abundance of different RNA types differed markedly across tissues (**Figure 2A**), with piRNAs being most enriched in the testis, consistent with their known germline-specific expression [47–49]. CA showed the highest percentage of fragments of rRNAs (rsRNAs), as reported previously [33, 50]. The most striking difference was in mitochondrial tRNAs (mt-tRNAs), with heart tissue exhibiting the highest proportion of mt-tRNA reads. This enrichment aligns with prior studies [2, 4, 51, 52] and reflects the heart’s high mitochondrial demand. At the level of specific small RNAs, OTTR-seq captured well-established tissue-specific miRNA expression patterns (**Figure S3**). For example, miR-122 and miR-192 were highly enriched in liver tissue [53], while miR-499 and miR-1a-1 were detected exclusively in the heart [54]. Likewise, miR-465 [55] and miR-34c [56] were most abundant in the testis, consistent with previous findings. Together, these results validated the sensitivity and accuracy of OTTR-seq in detecting tissue-specific small RNA expression.

**Figure 2.**
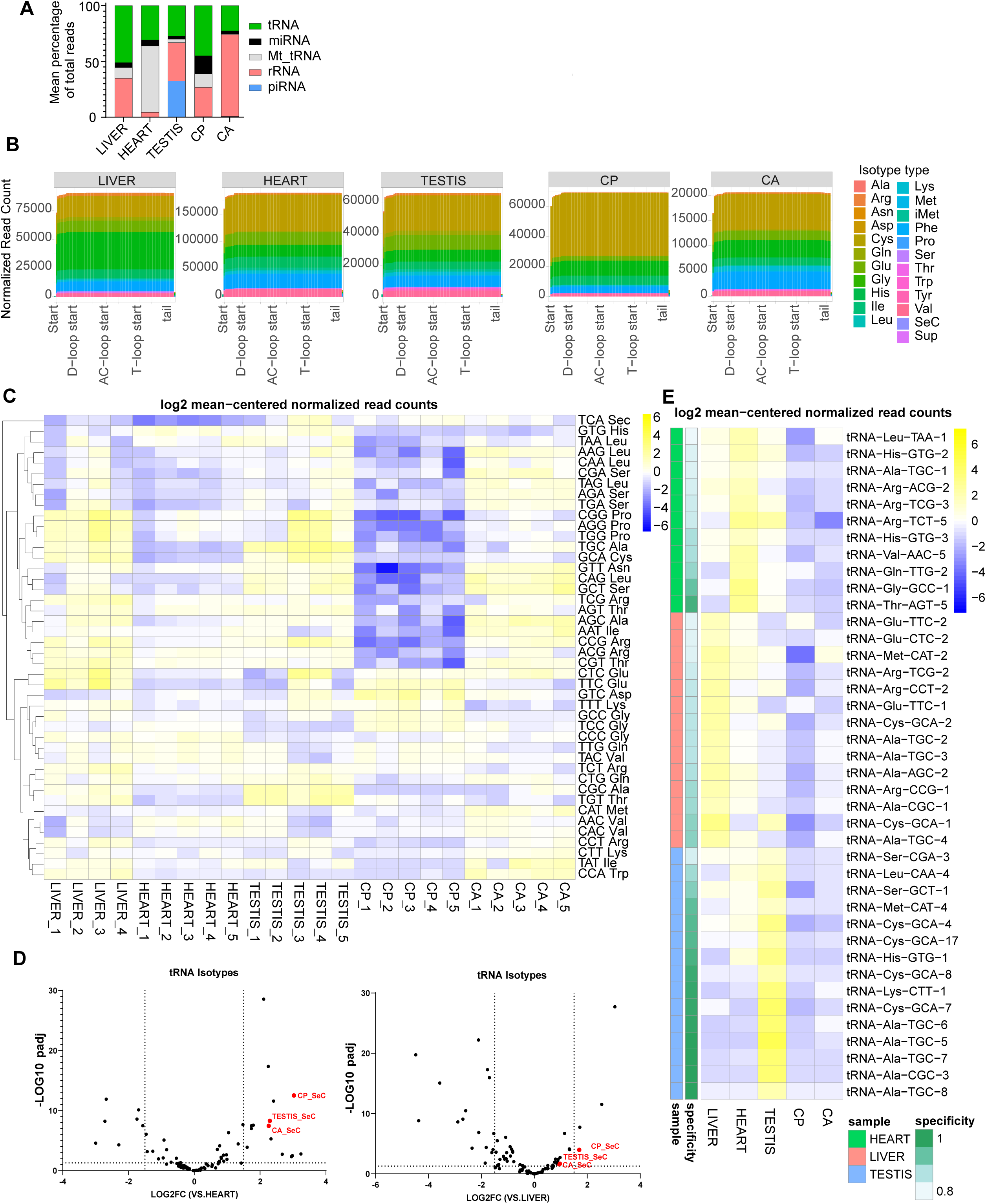
Tissue-specific tRNA abundance profiles. **A)** OTTR-seq analysis of the proportion of reads of different RNA classes across various tissues. Reads not mapping to annotated genes were excluded. **B)** Coverage plots showing normalized read counts for >65nts reads mapped to tRNA genes. The x-axis indicates nucleotide position, and the y-axis represents normalized read counts. Different tRNA isotypes are color-coded. **C)** Heatmap illustrating log2 mean-centered decoder-normalized read counts across all decoder types across various tissues (n=4-5 replicates per tissue). **D)** Volcano plot of differential expression of 21 isotypes in reproductive tissues compared to heart and liver samples, as determined by DESeq2. Differentially expressed tRNA-Sec in reproductive tissues is highlighted in red. **E**) Heatmap displaying log2 mean-centered tRNA-normalized read counts for the top 40 most sample-biased isodecoders along with their specificity score and maximally-abundant sample.

To investigate tissue-specific and diet-induced changes in tRNA abundance and modification, we focused on >65 nts reads mapping to tRNAs, which comprise full-length and near-full-length tRNAs, hereafter referred to as tRNA reads (**Table S1**). tRNA reads were summed by their respective amino acid decoding pool into 22 amino acid classes (tRNA isotypes), and read coverage plots were generated (**Figure 2B**). Next, tRNA reads that mapped to a single amino acid acceptor were classified into 47 anticodon-defined classes (tRNA isoacceptors), and their relative abundance across tissues was measured. This analysis revealed a differential abundance of numerous tRNA isoacceptors amongst these tissues (**Figure 2C**). Among the differentially expressed tRNAs, tRNA-Sec-TCA stood out for its significantly higher abundance in reproductive tissues compared to non-reproductive tissues (**Figure 2D**). tRNA-Sec-TCA is responsible for incorporating selenocysteine into selenoproteins. Selenoproteins have been implicated in maintaining sperm quality and male fertility, particularly by protecting sperm from oxidative stress during spermatogenesis [57–59]. The elevated levels of tRNA-Sec-TCA in the reproductive tissues may reflect a heightened demand for selenoprotein biosynthesis, supporting a specialized antioxidant environment crucial for maintaining sperm viability and function. To measure tissue-specific expression patterns of tRNA isodecoders, we calculated a tissue specificity index (TSI) [60]. This analysis revealed distinct tissue-biased abundance of several tRNA isodecoders amongst the set of analyzed samples, with the testis exhibiting the most and higher-scoring tissue-biased tRNAs (**Figure 2E**). These findings align with prior reports highlighting the testis as a tissue with uniquely high gene expression diversity and specificity [61, 62].

### Mature sperm are comprised of distinct nuclear and mitochondrial tRNAs

Following protamination, the paternal genome becomes transcriptionally inert and tightly compacted within the sperm nucleus [63, 64]. In the absence of cytosolic translational activity, it is expected that sperm lack full-length tRNAs. Surprisingly, we detected multiple tRNAs in sperm (**Figure 3A-B**), with the isotype composition of sperm tRNAs notably distinct from somatic and other reproductive tissues (**Figure 3C**). As observed in the testis and epididymis, tRNA-Sec levels were significantly higher in sperm relative to the liver and heart tissues (**Figure 3D)**. Within reproductive tissues, tRNA-Asp, tRNA-Glu, and tRNA-Gln were significantly less abundant in sperm compared to epididymal tissues (**Figure 3E**), suggesting a differential tRNA profile of the epididymis and sperm.

**Figure 3.**
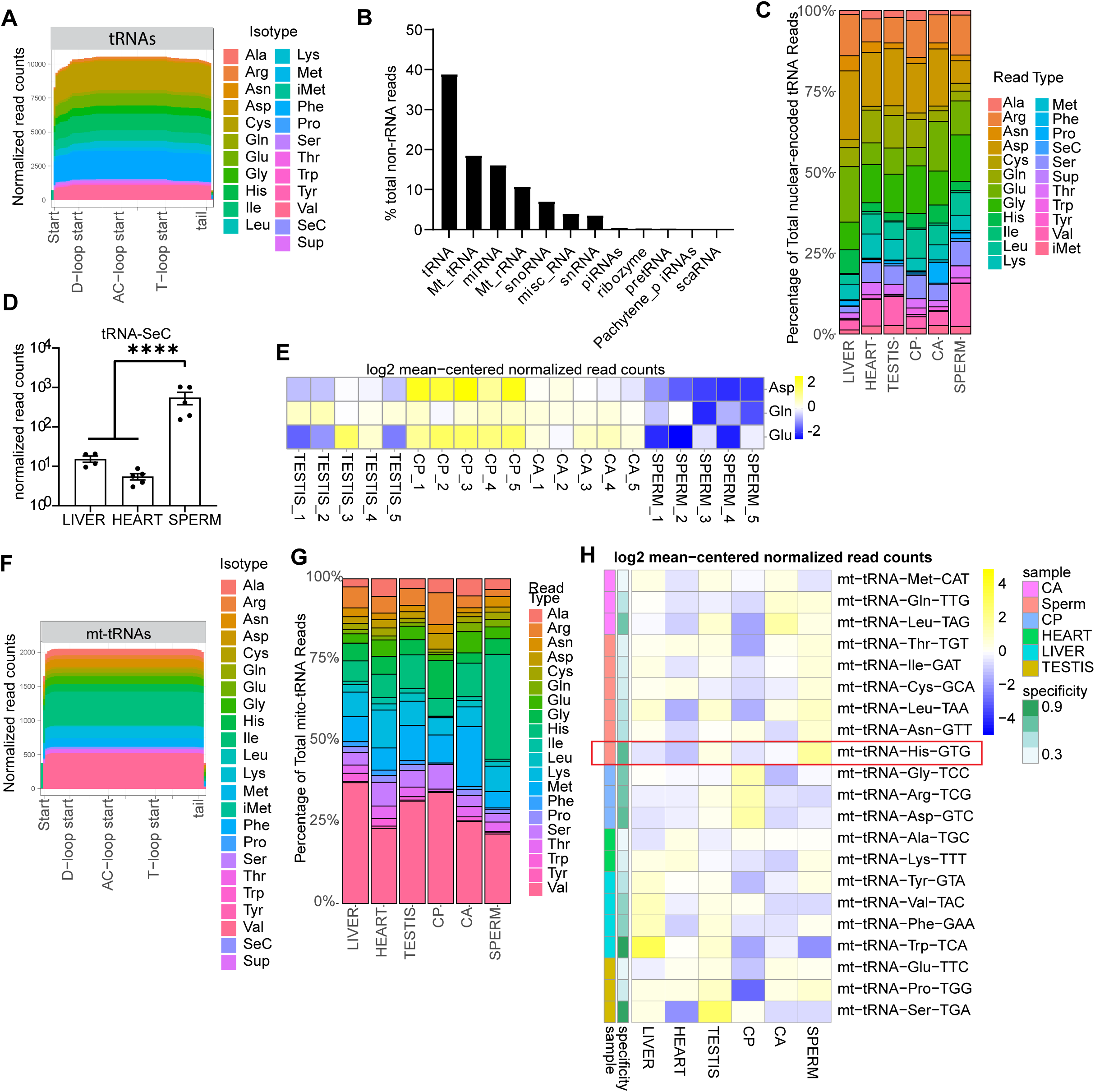
OTTR-seq analysis reveals distinct nuclear and mitochondrial tRNA profiles in sperm. **A)** Read coverage plots for >65 nts reads mapped to tRNA genes in sperm. The x-axis indicates nucleotide positions on tRNA genes, while the y-axis represents normalized read counts. Different tRNA isotypes are color-coded. **B)** Bar graph showing the percentage of reads mapped to each sRNA class in sperm. All reads (< and >65 nt reads) were included for this analysis. **C)** Bar graph showing the percentage of nuclear-encoded tRNA isotypes across tissues, with different isotypes color-coded. **D)** Bar graph of normalized read counts of tRNA-SeC in sperm, heart, and liver tissues. **E)** Heatmap displaying log2 mean-centered amino-normalized reads for acceptor types with differentially expressed tRNA isotypes in sperm compared to other reproductive tissues (testis, caput epididymis -CP, and cauda epididymis-CA). Statistical analysis was performed between sperm and each reproductive tissue with DEseq2 to identify commonly differentially expressed sperm-specific isotypes. Significant changes were determined using multiple t-tests with adjusted p-value <0.05. **F)** Sperm mitochondrial tRNA (mt-tRNAs) read coverage plots for >65 nts reads. **G)** Bar chart illustrating the percentages of mt-tRNA isotypes across all samples, with different isotypes color-coded. **H)** Heatmap showing log2 mean-centered normalized read counts of mt-tRNAs across all tissue samples with their specificity score and the maximally-abundant samples. mt-tRNA-His-GTG is highlighted in a red box as it shows sperm-biased expression.

We also detected full-length mt-tRNAs in sperm (**Figure 3B and F**), consistent with recent reports [32, 65]. While sperm and CA generally showed similar mt-tRNA expression patterns (**Figure 3H and S4**), mt-tRNA-His-GTG displayed sperm-biased expression (**Figure 3G-H**). Prior studies reported the active involvement of mitochondrial ribosomes in protein translation [66, 67]. High abundance of mt-tRNA-His-GTG in sperm suggests a potential role in regulating sperm mitochondrial biology. Histidine supplementation has been reported to protect against lead-induced reproductive toxicity by stabilizing mitochondrial function [68]. Higher levels of mt-tRNA-His-GTG could enhance mitochondrial protein synthesis, aiding cellular response to oxidative stress and preserving sperm function. Moreover, sperm mt-tRNAs can be modulated by diet in response to mitochondrial dysregulation and play a role in epigenetic inheritance of paternal dietary effects [65, 69]. Taken together, these data demonstrate the presence of mature, full-length nuclear and mitochondrial tRNAs in sperm and reveal a distinct expression profile that warrants further investigation for its potential role in sperm biology.

### Tissue-specific signatures of tRNA modifications revealed by mismatch profiling

The reverse transcriptase used in OTTR-seq can read through certain modified nucleotides more efficiently than other enzymes, which often stall at these sites. However, it frequently introduces errors by incorporating incorrect nucleotides or skipping bases at those positions. These misincorporation signatures in the sequencing reads can be used to infer the presence and location of specific tRNA modifications [70]. Indeed, we detected mismatches at known tRNA modification sites in nuclear-encoded tRNAs [30, 31, 42, 71], including position 9 (m1G), position 26 (m2,2G), position 34 (m3C and I), position 37 (m1G, m1I, and others), and position 58 (m1A) (**Figure 4A**). We also detected misincorporations at previously reported mt-tRNA modification sites (**Figure 4B**) [72]. Notably, the m¹A9 modification has been identified in most mt-tRNAs [72] and is crucial for proper tRNA folding [73]. Consistent with these reports, we detected mismatch signatures at position 9 in most mt-tRNAs (**Figure 4B**). We also detected misincorporation at positions 26, 32, and 37 in a subset of mt-tRNAs previously reported to be modified in humans (**Figure 4B and S5)** [72]. For instance, mt-tRNA-Pro and mt-tRNA-Leu, which harbor m¹G at position 37 in humans, showed clear mismatch signals in mouse tissues (**Figure S5C**). Similarly, mt-tRNA-Tyr, -Trp, and -Phe, known to carry ms²i⁶A at position 37, exhibited misincorporation at this site. At position 32, mismatches were observed exclusively in mt-tRNA-Ser and -Thr (**Figure S5B**), consistent with reported m³C modifications in humans. In contrast, position 26 showed broader misincorporation across mt-tRNA-Gln, -Glu, -Leu, and -Lys in our mouse data (**Figure S5A**), whereas human data identified m²G or m²,²G modifications primarily in mt-tRNA-Ala, -Glu, -Arg, and -Ile. These results suggest species-specific differences exist in the modification landscape, while many mt-tRNA modification sites are conserved.

**Figure 4.**
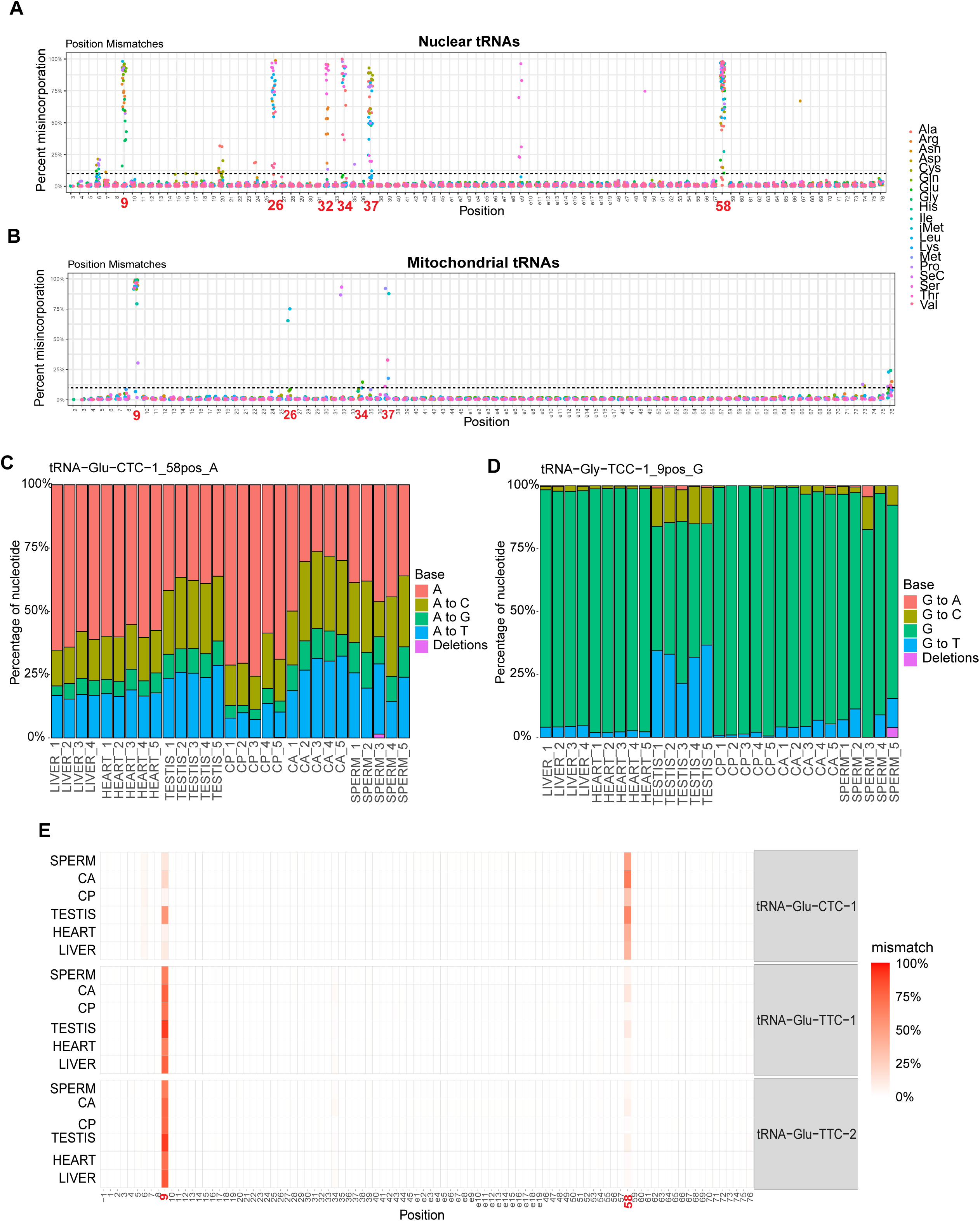
Tissue-specific misincorporation signatures of tRNA modifications. **A-B**) Plots showing misincorporation rates across all positions (x-axis) for nuclear-encoded tRNAs (A) and mitochondrial tRNAs (B), color-coded by isotype. Previously characterized modified positions are highlighted in red. Plots showing maximum misincorporation rates for all samples for each tRNA. **C**) Misincorporation plot showing mismatch rates at position 58 on tRNA-Glu-CTC-1 across all tissue types. The plot compares the percentage of each nucleotide or deleted (skipped) base detected at this position. As described in the key, A is the reference base, and A to C, A to G, A to T, or deletion are the misincorporation signatures. **D)** Misincorporation plot showing mismatch rates at position 9 on tRNA-Glu-TCC-1 across all tissue types. The plot compares the percentage of each nucleotide or deleted (skipped) base detected at this position. **E)** Heatmap showing average mismatch levels at each position for Glutamic acid tRNAs.

As the specific misincorporation pattern at modified bases depends on the sequence context, the type of RT enzyme used, and other influencing factors [74], absolute modification levels cannot be inferred solely from mismatch frequency. However, relative differences in base misincorporation across samples can be used to examine changes in modification status at specific positions. A comparison of misincorporation levels across tissues revealed 31 unique tRNA positions with differential misincorporation rates between tissues (**Table S2;** significance cut off used: p value <0.05, Mann-Whitney U-test). For example, an unmodified adenosine at position 58 is read correctly as “A,” whereas a m¹A modification typically results in a mismatch (A to T, A to C, or A to G) or deletion. We observed tissue-specific variation in mismatch signature at A58 (**Figure 4C and S6A-D**), including modest differences across isoacceptors such as tRNA-Val-CAC and tRNA-Val-TCA (**Figure S6A-B**). For tRNA-Glu, the misincorporation rate in tRNA-Glu-CTC-1 at A58 was significantly higher in testis and CA (**Figure 4C and E)**. Importantly, these differences were not driven by changes in the abundance of tRNA-Glu-CTC-1, as its expression levels did not correlate with the extent of m¹A58 modification across tissues (**Figure S6E**). In contrast, tRNA-Glu-TCT-1 and tRNA-Glu-TCT-2 were minimally modified at A58 and highly modified at position 9 (**Figure 4E**), revealing isoacceptor-specific modifications in this tRNA.

In addition to position 58A, we observed tissue-specific misincorporation signatures at other known modification sites, including position 9 (m¹G) and position 32 (m³C) (**Figure 4D-E and S6F**). For example, in tRNA-Glu, the mismatch rate at position 9G was markedly higher in testis compared to other tissues (**Figure 4E**). Similarly, for tRNA-Gly, we detected high levels of misincorporation at position 9 in the testis tissues amongst all sequenced isoacceptors (**Figure 4D and S7**). This pattern is consistent with previously reported high expression levels of the m¹G methyltransferase enzyme Trmt10a in the testis, suggesting a potential link between enzyme expression and tRNA modification at this site [75]. Although the functional significance of tissue-specific modification signatures is unclear, these modifications could regulate translation, tRNA stability, biogenesis of fragments, or interactions with other proteins in a tissue-specific manner.

To explore whether misincorporation levels, which reflect underlying tRNA modifications, are associated with the abundance of the corresponding tRNA molecules, we examined the correlation between mismatch rates and tRNA expression levels at frequently modified positions, including positions 9, 26, 32, 34, 37, and 58. This analysis revealed no consistent association between misincorporation frequency and tRNA abundance across tissues (**Figure S8**). However, we cannot rule out the possibility that such relationships may exist at the single-molecule level, requiring future studies employing higher-resolution techniques to examine modification status on individual tRNA transcripts.

### Low protein diet alters tRNA abundance and key modifications in the heart and liver

Next, we assessed the impact of dietary interventions on tRNA abundance and modifications. Comparing LP and HF diets to their corresponding controls showed that the high-fat diet induced minimal changes in tRNA profiles (**Figure S9A**). In contrast, the LP diet led to distinct alterations in liver and heart tissues at the level of isotypes and isoacceptors (**Figure 5A–D**). A subset of these changes in tRNA abundance likely reflect upstream alterations in specific isodecoders (**Figure 5E-F**). For example, the reduced abundance of tRNA-Val in LP heart tissue (**Figure 5B**) corresponded to the downregulation of tRNA-Val-CAC, tRNA-Val-AAC, and tRNA-Val-TAC isoacceptors (**Figure 5D**). Further dissection revealed specific isodecoders—tRNA-Val-CAC-2, tRNA-Val-AAC-1, tRNA-Val-AAC-5, and tRNA-Val-TAC-1—as contributors to these changes (**Figure 5F**).

**Figure 5.**
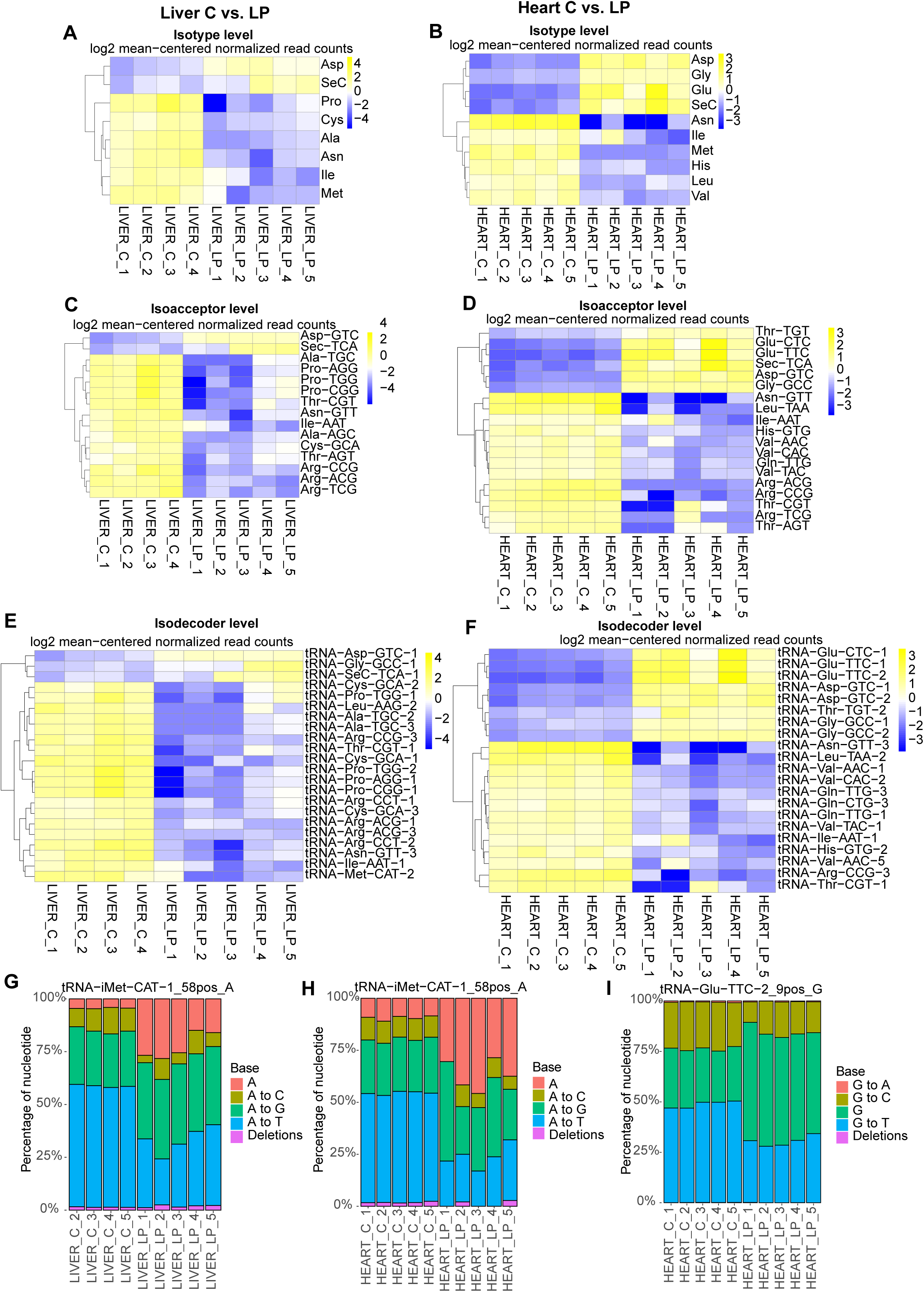
Impact of diet on tRNA abundance and modifications in the liver and heart. **A-B)** Heatmaps displaying log2 mean-centered acceptor-normalized read counts of significantly altered tRNA isotypes in LP liver (A) and LP heart (B) compared to corresponding control diet samples. **C-D)** Heatmaps showing log2 mean-centered decoder-normalized read counts of significantly altered tRNA isoacceptors in LP liver (C) and LP heart (D) compared to control diet samples. **E-F)** Heatmaps depicting log2 mean-centered tRNA-normalized read counts of significantly altered tRNA isodecoders in LP liver (E) and LP heart (F) compared to control diet samples. The DEseq2 cutoff criteria for significance were set to log2FC > 1.5 and an adjusted p-value < 0.05. **G-H)** Mismatch rates at position 58 of tRNA-iMet-CAT-1 in control and LP liver and heart samples, with nucleotide percentages represented by different colors and reference bases as A. **I)** Mismatch rates at position 9 of tRNA-Glu-TTC-2 in control and LP heart samples, with nucleotide percentages represented by different colors.

To identify tRNA modification changes in response to dietary challenges, we next analyzed misincorporation rates among tissues from the LP and HF groups. In response to dietary challenges, 17 unique tRNA positions exhibited differential misincorporation rates across tissues (**Table S2;** significance cut off used: p value <0.05, Mann-Whitney U-test). For example, in the liver tissues, tRNA-iMet-CAT-1 exhibited a lower misincorporation rate at A58 (reflecting lower m1A58 levels) in LP liver compared to the control group (**Figure 5G**). In the LP heart tissues, there was a significant decrease in the levels of misincorporation at A58 in tRNA-Glu-CTC-1, tRNA-Asp-GTC-2, tRNA-Gly-CCC-1, tRNA-Gly-GCC-1, tRNA-Gly-GCC-2, tRNA-Gly-TCC-1, as well as tRNA-iMet-CAT-1 (**Figure 5H and S9B)**. There was also a significant decrease in misincorporation rate at position 9 (likely due to reduced m1G modification) in the LP heart tissue in tRNA-Glu-TTC-1 (**Figure 5I**). Given that m1A58 modification in tRNA-Glu-CTC-1 is also regulated in a tissue-specific manner (**Figure 4D-E**), it is tempting to speculate that this site is highly dynamic and can be controlled by intrinsic and extrinsic factors. Moreover, we detected reduced m1A58 in initiator tRNA-iMet-CAT-1 in both tissues, suggesting a systemic effect on translation initiation [76–78]. m^1^A58 and m1G9 modifications contribute to stabilizing tRNA structure and, thereby, its stability [79–82]. Thus, a reduction in these modifications in response to dietary protein restriction may compromise tRNA structural stability, potentially affecting translation efficiency and fidelity.

To our surprise, the HF diet did not lead to significant changes in tRNA modifications in the OTTR-seq data, whereas LC-MS/MS detected robust differences. As noted above, for the OTTR-seq experiments, we used a dedicated HF diet control (C_HF), which differs from the HF diet only in the fat content. In contrast, the LC-MS/MS experiments used a common control diet (C) for both LP and HF groups. Interestingly, tissues from mice fed the C_HF diet showed lower modification levels at specific tRNA positions compared to those fed the C diet (**Figure S10**), indicating that the two control diets differ in their impact on tRNA modifications. A comparison of their compositions revealed substantial differences in both macronutrient profiles and ingredient sources: the C diet is soybean oil–based and rich in polyunsaturated fats, whereas the C_HF diet contains lard, leading to higher levels of saturated and monounsaturated fats. These compositional differences may independently influence tRNA abundance and modification patterns, underscoring the importance of careful selection and reporting of control diets in dietary intervention studies.

### Diet modulates the abundance of specific tRNAs in reproductive tissues with minimal effects on modifications

We examined how dietary interventions impact tRNA abundance and modifications across reproductive tract tissues. Unlike the robust responses seen in liver and heart, reproductive tissues generally exhibited minimal tRNA abundance changes under both HF and LP diets, except for CP (**Figure 6A-D)**. At the isodecoder level, four tRNAs were consistently upregulated in both LP and HF CP tissues: tRNA-Glu-CTC-1, tRNA-Glu-TTC-1, tRNA-Glu-TTC-2, and tRNA-Pro-TGG-2 (**Figure 6A-B**). These alterations at the isodecoders level contribute to an increase in tRNA-Glu-TTC and tRNA-Glu-CTC isoacceptors levels (**Figure 6C-D**) and an overall increase in tRNA-Glu isotype levels in both LP and HF diet groups relative to controls (**Figure 6E**). These shared changes suggest activation of common molecular pathways in response to both nutritional challenges. In addition, tRNA-Asp-GTC-1 and tRNA-Asp-GTC-2 were upregulated in CA of LP mice (**Figure 6F**), and tRNA-Glu-CTC-1 levels increased in sperm of the HF group (**Figure 6G**). However, these isodecoder-level changes did not contribute to isotype-level shifts, indicating possible translation-independent roles or other specialized functions. Notably, no significant changes in tRNA abundance were detected in the testis under either dietary challenge. These findings highlight the CP as the most diet-sensitive reproductive tissue in our analysis. Consistent with LC-MS/MS analysis, we did not detect significant changes in tRNA modifications in response to dietary challenges in the reproductive tissues.

**Figure 6.**
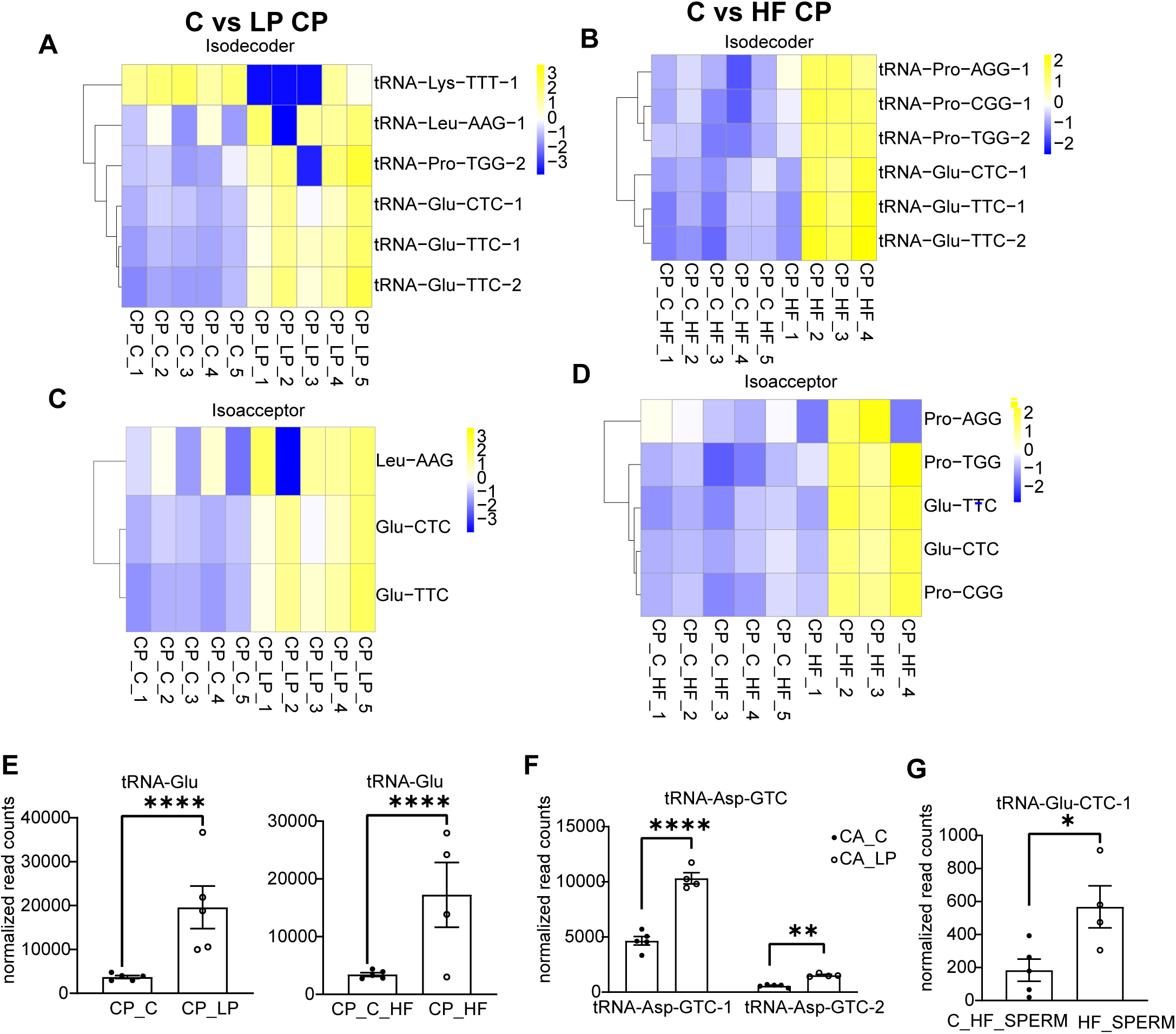
Impact of diet on tRNA levels and modifications in reproductive tissues. **A-B)** Heatmaps showing log2 mean-centered tRNA-normalized read counts of significantly altered tRNA isodecoders in caput epididymis (CP) under low-protein (LP) diet (A) and high-fat (HF) diet (B). **C-D)** Heatmaps displaying log2 mean-centered tRNA-normalized read counts of differentially expressed tRNA isoacceptors in CP under LP diet (C) and HF diet (D). **E)** Bar plot shows normalized read counts of significantly differentially expressed tRNA-Glu in CP under LP diet and HF diet. **F)** Bar plot shows normalized read counts of two differentially expressed isodecoders, tRNA-Asp-GTC-1 and tRNA-Asp-GTC-2, in the cauda epididymis (CA) under the LP diet. **G)** Bar plot displaying normalized read counts of tRNA-Glu-CTC-1, the only differentially expressed isodecoder in sperm under the HF diet.

### Dietary conditions alter mitochondrial-tRNA abundance and modifications

Given the observed alterations in nuclear-encoded tRNAs, we next investigated whether mt-tRNA levels and modifications are also affected by dietary changes. The mitochondrial genome encodes 22 distinct tRNAs, each representing a unique isotype [83]. Similar to nuclear-encoded tRNAs, liver and heart tissues showed the most pronounced diet-induced changes in mt-tRNA abundance in response to the LP diet (**Figure 7A–B**). Notably, several tRNAs—including mt-tRNA-Thr-TGT, mt-tRNA-Leu-TAG, and mt-tRNA-Tyr-GTA—were consistently downregulated in both LP liver and LP heart, suggesting shared metabolic responses to protein restriction (as observed for a subset of nuclear-encoded tRNAs). In contrast to liver and heart, reproductive tissues exhibited more limited changes in mt-tRNA expression in response to dietary interventions. Specifically, mt-tRNA-Ser-TGA was downregulated in HF testis, mt-tRNA-Asp-GTC was downregulated in the LP CP, and mt-tRNA-Trp-TCA was upregulated in HF sperm (**Figure 7C**). The functional significance of these alterations remains to be investigated. Intriguingly, while the nuclear-encoded tRNA-Val-TAC was downregulated in the LP heart (**Figure 5B, D, and F**), mt-tRNA-Val-TAC was upregulated in this tissue (**Figure 7B and S11A**). These data suggest differential control of tRNA levels for pairs of isoacceptors partitioned between mitochondria and the nucleus, as observed previously in response to cellular stress [9]. Differential abundance of the same isoacceptors in the cytosol and mitochondria may enable distinct translational regulation of protein restriction response in each compartment.

**Figure 7.**
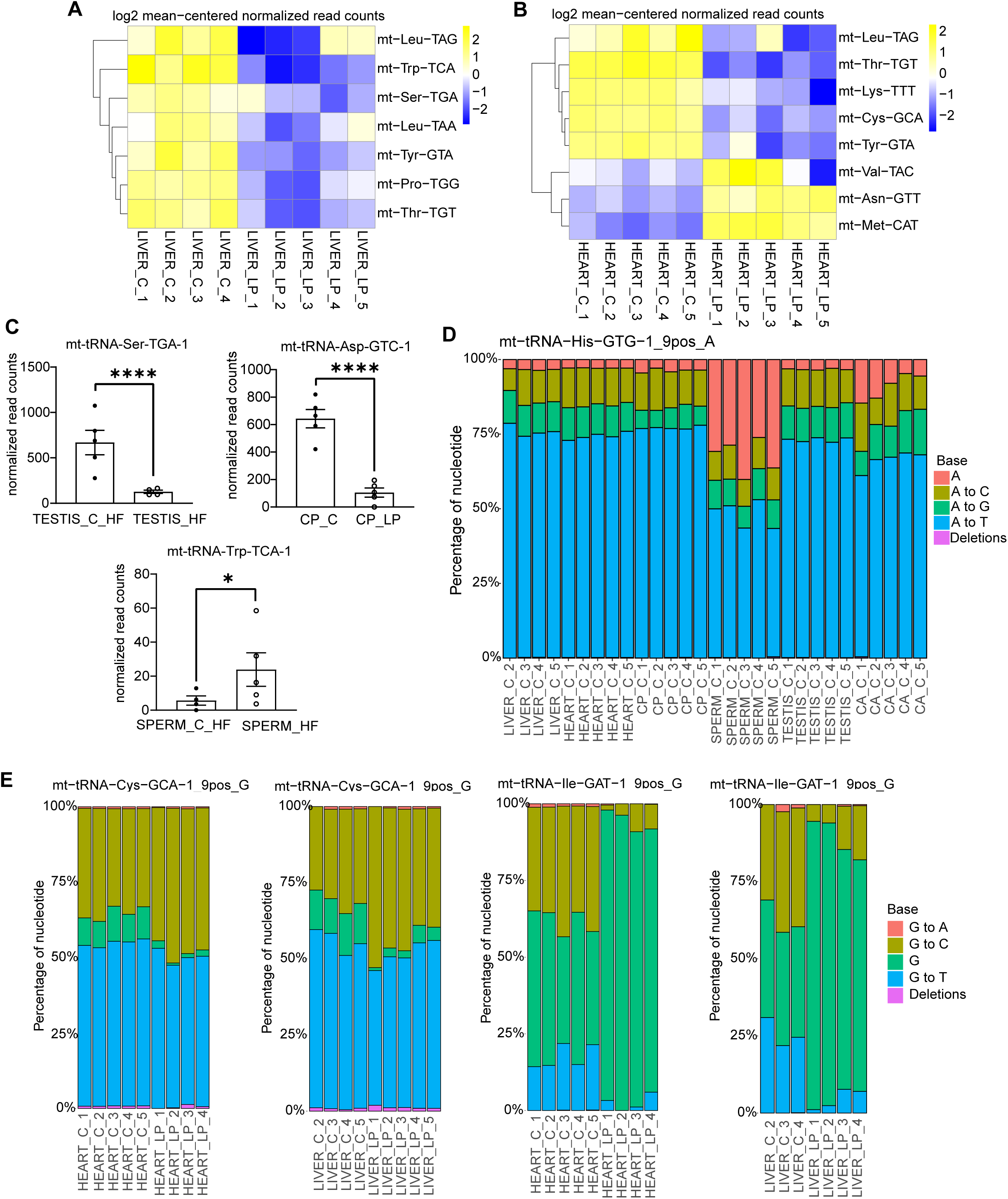
Diet-induced changes in mitochondrial-tRNA abundance and modifications. **A-B)** Heatmaps showing log2 mean-centered mt-tRNA normalized read counts of mitochondrial tRNA isodecoders differentially expressed between control liver (LIVER_C) and low-protein liver (LIVER_LP) samples (A) and between control heart (HEART_C) and low-protein heart (HEART_LP) samples (B). **C)** Bar plot showing normalized read counts of differentially expressed mitochondrial tRNAs in high-fat (HF) sperm and testis, and low-protein (LP) caput epididymis (CP). **D)** Mismatch plot illustrating the percentage of each nucleotide detected at position 9 of mt-tRNA-His-GTG-1 across all tissue types, including sperm. Colors distinguish different nucleotides. Liver_C_1 was excluded due to insufficient coverage (<20 reads at position 9 of mt-tRNA-His-GTG-1). **E)** The mismatch incorporation rates (y-axis) for position 9 are displayed for two mitochondrial tRNAs, mt-tRNA-Cys-GCA and mt-tRNA-Ile-GAT, comparing control and LP diet conditions in heart and liver tissues. Heart_LP_5 samples is omitted from the plots because it had fewer than 20 reads covering position 9. Similarly, due to insufficient read coverage at position 9 for the mt-tRNA, Liver_C_1 is excluded from both plots for mt-tRNA-Cys-GCA and mt-tRNA-Ile-GAT, and Liver_LP5 is excluded from the mt-tRNA-Ile-GAT plot. In all cases, at least 3 replicates are analyzed.

We identified 9 unique mt-tRNA positions that exhibited differential misincorporation rates across tissues, and 6 that displayed differential misincorporation rates in response to dietary challenges (**Table S3;** significance cut off used: p value <0.05, Mann-Whitney U-test). For example, sperm exhibited the lowest levels of m¹A at position 9 in mt-tRNA-His-GTG compared to other tissues (**Figure 7D**). Additionally, reproductive tract tissues—including CP, testis, and sperm—showed higher levels of m¹G at position 26 in mt-tRNA-Lys-TTT compared to somatic tissues such as liver and heart (**Figure S11B**). Similar to nuclear-encoded tRNAs, diet-induced changes in mt-tRNA modifications were primarily observed in the liver and heart tissues in response to the LP diet (**Figure 7E, S11C**). For instance, the LP diet led to increased m¹G at position 9 in mt-tRNA-Cys-GCA and decreased m¹G9 levels in mt-tRNA-Ile-GAT in both tissues (**Figure 7E**). Reproductive tissues were largely unaffected by dietary intervention, with one exception: mt-tRNA-Gln-TTG exhibited elevated m¹G9 levels in LP-fed sperm (**Figure S11D**). These findings indicate that while dietary composition modulates mitochondrial tRNA dynamics, these effects are tissue-specific and are most pronounced in somatic tissues, with limited impact on reproductive tissues.

## Discussion

Our findings demonstrate that tRNA modifications and abundance are regulated in a tissue-specific and diet-responsive manner, adding to the growing recognition of tRNA molecules as dynamic regulators of cellular function.

Here, by comparing tRNA abundance in the male reproductive tract to heart and liver tissues, we comprehensively examine tRNA abundance in the male reproductive tissues and add to the growing appreciation of the unique profile of tRNAs in these tissues [32, 36]. We identified tRNA-SeC-TCA being enriched in reproductive tissues, which may be related to the role of selenoproteins in protecting sperm from oxidative stress [57, 59]. Moreover, amongst the tissues examined, the testis showed the highest number of tissue-biased tRNA isodecoders, including expression of brain-specific isodecoders of tRNA-Alanine (tRNA-Ala-TGC-5, tRNA-Ala-TGC-6, and tRNA-Ala-TGC-7) [2]. These findings align with prior observations of molecular and biochemical similarities between the brain and testis [84, 85]. Furthermore, OTTR-seq detected full-length nuclear-encoded tRNAs in sperm and high levels of mt-tRNA-His-GTG. A higher abundance of mt-tRNA-His in sperm may be attributed to the enrichment of mitochondria in sperm, although the specific function of this tRNA remains to be deciphered.

Under LP and HF dietary challenge, liver and heart tissues showed the most robust changes in tRNA abundance, with limited alterations in reproductive tissues, suggesting that somatic tissues are more sensitive to nutrient composition. These differences likely reflect the distinct metabolic demands. For example, muscle-rich tissues like the heart rely heavily on proteins for repair and growth. Thus, heart tissue is particularly sensitive to dietary protein intake [86, 87]. Conversely, the liver, central to lipid metabolism, is particularly affected by high-fat intake and associated conditions such as non-alcoholic fatty liver disease [88, 89]. Finally, although limited, significant alterations were detected in specific tRNAs in the epididymis and sperm in response to different diets. A more comprehensive investigation is needed to delve into the biological consequences of these alterations on male reproduction.

In addition to profiling tissue-specific and diet-sensitive tRNA abundance, we detected potential tissue-enriched tRNA modifications and the dietary effects on those modifications using LC-MS/MS and mismatch signatures in OTTR-seq reads. For instance, LC-MS/MS data revealed that gal-Q modification is higher in the epididymis relative to the liver. The testis tissues showed increased levels of m1A58 and m1G9 compared to other tissues in an isodecoder-specific manner. Moreover, the liver and heart tissues exhibited a decrease in tRNA modification levels at position 58A in nuclear-encoded tRNA and 9A in mitochondrial tRNA when mice were subjected to a low-protein diet. These specific modifications are known to play important roles in tRNA stability and secondary structure maintenance [80, 81]. As shown in the schematic in **Figure 1H**, many of the RNA modifications regulated in a tissue-specific manner or altered by LP and HF diets are located around the anticodon region, emphasizing their functional significance in translation regulation. Notably, modifications such as m⁶t⁶A, i⁶A, and galQ are well-characterized for enhancing tRNA decoding efficiency and translational fidelity [90–92]. The observed alterations in these modifications in response to dietary stress suggest a potential mechanism by which tRNAs modulate translational output to adapt to changes in nutrient availability or metabolic demand [93, 94].

A limitation of our study is that cDNA-based tRNA sequencing methods rely on amplification steps and may not reflect the true abundance of tRNA molecules, particularly those with extensive modifications. Direct RNA sequencing platforms such as Oxford Nanopore Technologies (ONT) offer promise to overcome this, having been successfully applied to yeast and E. coli tRNA pools [31, 95]. However, ONT-based methods are still developing for highly modified and structurally complex mammalian tRNAs, and further optimization is needed for routine application in higher organisms.

In conclusion, our study reveals that tRNA abundance and modifications are intricately shaped by both tissue context and dietary composition, offering new insight into how cells may fine-tune their translational machinery in response to environmental and physiological cues. These findings lay a foundation for future research into the functional consequences of tRNA dynamics in health, disease, and inheritance.

## Methods

### Mouse husbandry

Wild-type FVB/NJ mice were used in the study. All animal care and use procedures were approved by the University of California, Santa Cruz Institutional Animal Care and Use Committee. Mice were group-housed (maximum of 5 per cage) with a 12-hour light-dark cycle (lights off at 6 pm) and free access to food and water *ad libitum*. Animals were raised on one of three diets – defined Control diet (BioServ AIN-93g, cat# F3156) or a low protein diet based on AIN-93g (10% of protein rather than 18%, remaining mass made up with sucrose, BioServ, cat# S3156) or high fat diet (60% of fat calories rather than 16%, BioServ, cat# S3282) for the data in Figure 1 and Supplementary Figure 1. For the rest of the data for OTTR-seq, mice were fed on the above diets with one additional diet (7.2% fat calories, BioServ, cat# S4031) as a control diet for the high-fat diet. Littermates were split into the treatment diet and its corresponding control diet.

### Mouse tissue collection and RNA isolation

All mouse tissue samples were isolated from wild-type FVB/NJ mice using procedures approved by the University of California, Santa Cruz Institutional Animal Care and Use Committee. Samples were rapidly frozen in liquid nitrogen and stored at −80 °C until use. Then, 1 mL of TRIzol (Invitrogen, cat # 15596026) was added per 100 mg of dissected whole tissue, and samples were homogenized in TRIzol buffer using a homogenizer until the suspension was completely homogeneous. We performed homogenization using the following settings: medium speed for 3-5 cycles (10 seconds per cycle) at 4°C. Cell debris was removed by centrifugation at 12000 RCF for 10 minutes, followed by phase separation and isopropanol precipitation (as described above). RNA pellets were resuspended in nuclease-free water and stored at −80°C until use.

### Sperm preparation and RNA isolation

Epididymis dissection, sperm isolation, and RNA isolation were performed following previous protocol [20]. Caput and cauda epididymis tissues were dissected out from an adult mouse and transferred to a 35mm dish containing 1ml of pre-warmed Whitten’s Media (100 mM NaCl, 4.7 mM KCl, 1.2 mM KH2PO4, 1.2 mM MgSO4, 5.5 mM Glucose, 1 mM Pyruvic acid, 4.8 mM Lactic acid (hemicalcium), and HEPES 20 mM). Sperm were obtained by first making two incisions at the caput or cauda then gently squeezing the tissue to release fluid. After a 15-minute incubation at 37°C, only cauda sperm-containing media were transferred to a new tube for an additional 15-minute incubation. Sperm-free epididymis tissues were frozen directly in liquid nitrogen. After a 30-minute incubation at 37°C, sperm were collected by centrifugation at 2000 x g for 2 minutes, washed with 1X PBS, and then washed with lysis buffer for 10 minutes on ice to remove somatic cell contamination. The sperm sample was finally washed with 1X PBS and pelleted after centrifuged at 2000 × g. The resulting sperm pellets were then subjected to RNA extraction. For sperm RNA extraction, samples are first thawed if flash frozen previously. Otherwise, adjust the total volume to 60 µl with filtered water. Then add 33.3 µl of lysis buffer (6.4 M Guanidine HCl, 5% Tween 20, 5% Triton, 120 mM EDTA, and 120 mM Tris pH 8.0), 3.3 µl ProteinaseK (>600 mAU/ml, Qiagen, cat# 19131), and 3.3 µl 0.1M DTT to the sample. The sperm pellet was disturbed physically using a pipette tip, then incubated, with shaking, at 60°C for 15 minutes on an Eppendorf thermomixer. One volume of water (100 µl) was then added, and the samples were subjected to phase separation using TRI Reagent (Invitrogen) and 40 µl BCP (1-bromo-2 chloropropane, Sigma-Aldrich). Samples were vortexed for 5 minutes to ensure complete breakdown of the sperm, followed by centrifugation at 12,000 RCF for 4 minutes. The aqueous phase was then removed and transferred to a low-binding RNase/DNase-free microcentrifuge tube, followed by the addition of 20 µg of glycoblue (Invitrogen) and 1 volume of Isopropanol. The RNA was then precipitated for 30 minutes or greater at -20°C, followed by centrifugation at 18,000 RCF for 30 minutes at 4°C, and one wash with 70% cold ethanol followed by centrifugation at 12,000 RCF for 10 minutes at 4°C. Finally, the RNA was reconstituted in nuclease-free water.

### LC-MS/MS based RNA modification analysis

For LC-MS/MS nucleoside analysis, RNA samples were digested to nucleosides using a previously described method [96]. Briefly, the RNA samples were denatured at 95 °C for 3 minutes and immediately cooled. Samples were incubated for 2 hours with nuclease P1 (0.1 U/μg RNA) (Sigma-Aldrich) in 0.01 M ammonium acetate. The samples were then incubated for 2 hours at 37°C with snake venom phosphodiesterase (1.2×10^−4^U/μg RNA) and bacterial alkaline phosphatase (0.1U/μg RNA) (Worthington Biochemical corporation) in 1/10 volume of 1 M ammonium bicarbonate. A TSQ Quantiva Triple Quadrupole (Thermo Fisher Scientific) coupled to a Vanquish Flex Quaternary (Thermo Fisher Scientific) with an Acquity UPLC HSS T3, 1.8 μm, 1.0 mm x 100 mm column (Waters) at 30°C was used for analysis. The nucleosides were eluted with 5.3 mM ammonium acetate (pH = 4.5)(mobile phase A) and 5.3 mM ammonium acetate (pH = 4.5) in 40% acetonitrile (mobile phase B) using a previously described LC elution method [97]. A selected reaction monitoring method was used to quantify RNA modifications in samples with predetermined SRM transitions. Analysis was done in positive polarity with an H-ESI source. The Q1 and Q3 resolution was 0.7, and a Dwell Time of 300 ms. For collision induced dissociation (CID), nitrogen gas was used at 1.5 mTorr. The global parameters used were sheath gas, auxiliary gas, and sweep gas of 35, 10, and 0 arbitrary units, respectively; ion transfer tube temperature of 325 °C; vaporizer temperature of 275 °C; and spray voltage of 3.8 kV. Data acquisition and processing for relative quantification was done with Xcalibur 4.2 (Thermo Fisher Scientific). Modification levels were quantified by normalizing each modified base (e.g., m¹G) to the level of its corresponding unmodified canonical base (e.g., G).

### PNK and deacylation of RNA sample for sequencing

For PNK treatment, 1ug total RNA was treated with 5U PNK (NEB, cat# M0201S) in 5x PNK buffer (350 mM Tris-HCl pH 6.5, 50 mM MgCl2, 5 mM DTT) with 10U RNase inhibitor (Invitrogen, cat# AM2694). Reaction incubated at 37°C for 30 minutes in a thermocycler. To deactivate T4 PNK, 0.5 M EDTA was added, and samples were returned to incubation at 65°C for 15 minutes to deactivate T4 PNK. To deacylated acylated-tRNA, 3.5 µL of 100 mM Na2B4O7·10H2O (Sodium Borate; Sodium Tetraborate, Borax) was added to raise pH to ∼9 and incubated at 45°C for 30 minutes. Next, RNA purification was performed using phase separation followed by isopropanol precipitation, as described previously.

### Ordered two-template relay sequencing (OTTRseq)

OTTR was performed as previously described [98]. Briefly, input RNA was labelled at the 3’ end by incubation in buffer containing ddATP for 90 minutes in 30°C, followed by an addition of ddGTP and another incubation at 30°C for 30 minutes. The reaction was stopped by incubating at 65°C for 5 minutes, followed by the addition of 5 mM MgCl2 and 0.5 units of shrimp alkaline phosphatase (NEB, cat# M0371S) at 37°C for 15 minutes, then stopped by the addition of 5 mM EGTA, and incubated at 65°C for 5 minutes. Samples were then incubated in templated cDNA synthesis buffer, adaptors, and dNTPs at 37°C for 20 minutes, followed by heat inactivation at 65°C for 5 minutes and RNase A/H treatment. cDNA was size selected on a 10% PAGE urea gel to minimize adaptor dimers. Size selected cDNA was PCR amplified for 12 cycles with Q5 high fidelity polymerase (NEB, cat# M0491S). The final PCR product was cleaned up with AMPure XP beads (Beckman, cat # A63881) and finally separated by 6% TBE gel to remove adaptor dimers. The desired product was excised from the gel and eluted in 400ul elution buffer (300 mM NaCl, 10 mM Tris pH 8, 1 mM EDTA) overnight at -20, followed by isopropanol precipitation and resuspension in 9µl nuclease-free H2O.

### Small RNA sequencing analysis

The small RNA sequencing data subjected to tRAX analysis pipeline. Briefly, sequencing reads were trimmed with cutadapt with adapter “--no-indels -O 15 -m 2 -a GATCGGAAGAGCACACGTCT” and further trimming the final TRPT base with “cutadapt -u -1” and removal of the UMI with “cutadapt -u 7.” Sequencing analysis was done using a modified version of the tRAX analytical analysis tool [99] with default parameters except for a minimum non-tRNA read size of 16 and using the Ensembl gene set [100], the mm10 piRNA gene set from piPipes [101], and ribosomal RNA repeats from UCSC repeatmasker [102] as the gene set. In tRAX, reads were mapped to the mouse mm10 genome combined with tRNA sequences taken from gtRNAdb [103]. All ribosomal repeat sequences were taken from the UCSC genome browser repeatmasker track for ribosomal RNAs. To create a sequence database for mature tRNA sequences, introns were removed, CCA tails were added, and “G” base was added to the start of histidine tRNAs. tRNA reads were defined as any reads whose best mapping includes a mature tRNA sequence. The tRAX pipeline uses bowtie2 with options "-k 100 --very-sensitive -- ignore-quals --np 5 –very-sensitive," and from that mapping extracts all best mappings from those results with exceptions for tRNA reads. For reads that mapped best to tRNAs, the non-tRNA reads were removed to prevent reads from redundantly mapping both to the mature sequence and the genome locus.. Read mappings were processed to ensure a single primary mapping for each read was marked and only this primary mapping was used to calculate percentages of total reads for gene types and acceptor types to prevent double-counting of gene types and lengths. Notably, epididymis tissues showed fewer full-length tRNA reads than the testis, heart, and liver. As all samples were treated the same way, the epididymis likely has a higher abundance of ribonucleases.

Normalized read counts, adjusted p-values and log2-fold change were calculated using DESeq2 [104] with default parameters as a component of the tRAX pipeline. Counts for isotypes, acceptor types, and individual decoders were analyzed and normalized separately within each category in order to calculate relative change. Plots were generated with ggplot2 and Prism. Mismatch U-test p-values were calculated by using U-tests to test for significance by comparing if all mismatch rates of a sample were greater than another, with 10% added to all samples to increase the stringency of the test. All mismatch tests excluded any samples with under 20 reads.

## Acknowledgements

We thank Jonathan Howard and Aidan Manning for training us on DM-tRNA-Seq and OTTR-seq library preparation protocols. We thank all the members of the Sharma lab for their helpful discussions on this work. This work was supported by NIH grant 1DP2AG066622-01 awarded to US and NIH GM058843 to PAL.

## Supplementary Material

### Supplementary Tables

**Table S1:** Total read counts for >65 nts transcripts. These raw read counts were normalized as part of the DESeq2 analysis pipeline and used for downstream data analyses.

**Table S2:** Table of mismatch rates comparison for each position in each nuclear tRNA for bases with >20 reads of coverage and tRNA base positions <3 removed. U-test values of mismatch rates in both directions calculated with values shifted by a 10% threshold to detect substantial changes.

**Table S3:** Table of mismatch rates comparison for each position in each mitochondrial tRNA for bases with >20 reads of coverage and tRNA base positions <3 removed. U-test values of mismatch rates in both directions calculated with values shifted by a 10% threshold to detect substantial changes.

### Supplementary Figures

**Figure S1. A)** Experimental design: 16–40 nts and 70–90 nts RNAs were size-selected from total RNA using polyacrylamide gel electrophoresis and processed for LC-MS/MS analysis to detect nucleotide modifications. **B)** Bioanalyzer traces showing the size distribution of the RNA samples used for LC-MS/MS. **C)** Levels of RNA modifications detected in the control epididymis (Epi) and liver tissues by LC-MS/MS, normalized to the level of canonical base (n=6 samples per tissue). **D)** DM-tRNA sequencing of 70-90 nts RNA samples used for LC-MS/MS. The bar graph illustrates the RNA composition on these samples, showing over 90% of reads corresponded to tRNAs, indicating the high purity of the size-selected tRNA samples. **E)** Nuclear tRNA reads from (D) were categorized by isotype, and the percentage of reads from each isotype was compared between liver and epididymis tissues to detect tissue-specific differences. **F)** Normalized read counts of tRNA-Tyr from DM-tRNA-seq of epididymis and liver tissues under control diet. **G)** Body weights of mice in the control (C), low-protein (LP), and high-fat (HF) diet groups were measured at the time of tissue collection. Mice in the HF diet group showed a significant increase in body weight compared to the control group. Statistical analysis was performed using the Mann-Whitney t-test, p-value < 0.0001.

**Figure S2. A)** Schematic of the experimental setup for the OTTR-seq experiment. Three-week-old male mice were fed one of four diets for six months: low-protein (LP), control for low-protein (C), high-fat (HF), or control for high-fat (C_HF). Tissues including liver, heart, caput epididymis (CP), cauda epididymis (CA), testis, and sperm were collected for downstream analyses. **B**) Histogram of read lengths for all mapped reads in all samples, with reads mapping to nuclear-tRNAs separated from other reads. The tRNA-mapped reads show a prominent 75 nt peak (full-length tRNAs) and shorter ∼35 nt fragments. **C**) Principal component analysis of the first two principal components of normalized read counts.

**Figure S3.** Normalized read counts of reported tissue-specific miRNAs.

**Figure S4.** Scatter plot illustrating the Pearson correlation between tRNA expression in sperm and CA (A), CP(B), or testis (C). A linear regression line is included, with the R-squared value indicating the strength of the correlation.

**Figure S5.** Violin plots showing misincorporation rate at position 26 (A), 32 (B), and 37 (C) of mt-tRNAs by acceptor type.

**Figure S6. A-D)** Mismatch incorporation rates (y-axis) showing the percentage of each nucleotide detected at position 58 for various tRNAs: tRNA-Val-CAC-2 (A), tRNA-Val-TAC-1 (B), tRNA-iMet-CAT-1 (C), and tRNA-Asp-GTC-2 (D), across all tissues. Sample CP_2 is omitted from (B) as insufficient reads (< 20) were detected for this tRNA at position 58. **E)** Normalized counts of reads mapped to tRNA-Glu-CTC-1 across the heart, caput epididymis (CP), and cauda epididymis (CA). The changes in abundance of this tRNA do not correlate with changes in misincorporation rate at position 58, as shown in Figure 4C. **F**) Misincorporation plot showing mismatch rates at position 32 on tRNA-Arg-CCT-4 across all tissues, with colors indicating different nucleotides. No reads were detected for this position in the following samples: CP_2, CP_5, CA_1, and CA_2. Sperm samples show the highest misincorporation rate at this position.

**Figure S7**. Heatmap showing average mismatch levels at each position for Glycine tRNAs.

**Figure S8. A-F**) Dot plots comparing mismatch rates to normalized read counts for specific tRNA positions. No correlation was observed for highly modified tRNA positions, such as 9, 26, 37, and 58. Positions with fewer than 20 reads were excluded from analysis.

**Figure S9. A)** Volcano plots showing significantly altered tRNA levels in HF liver (left) and HF heart (right). Differentially expressed tRNAs are highlighted, with significance determined by adjusted p-values. **B)** Heatmap showing average mismatch levels at each position for selected tRNAs showing less misincorporation at position 58 in heart low-protein samples, suggesting lower m1A levels.

**Figure S10. A-D**) Mismatch rates at position 58 of tRNA-Gly-TCC-1, tRNA-Gly-GCC-2, tRNA-Gly-GCC-1, and tRNA-Glu-CTC-1 in heart samples under all four diets. Heart_C_HF_4 was excluded from analyses A-C due to insufficient coverage (<20 reads at position 58). As shown, C_HF samples have lower misincorporation rates at this position compared to the C diet.

**Figure S11. A)** Stacked bar chart showing the percentage of mitochondrial tRNA isotypes in heart_C, heart_LP, CP_C, and CP_LP samples, with different colors representing individual mitochondrial tRNAs. As shown here, the overall distribution of tRNA isotypes is modulated by a low-protein diet in these tissues. **B)** Mismatch plots displaying the percentage of each nucleotide detected at position 26 on mt-tRNA-Lys-TTT-1 across all tissue types. Sperm and testis samples show the highest misincorporation rate at this position. **C-D)** Mismatch plots showing nucleotide percentage at key positions: mt-tRNA-Pro-TGG-1 (position 37), mt-tRNA-Leu-TAA-1 (position 26), and mt-tRNA-Thr-TGT-1 (position 32) in heart_C versus heart_LP (C) and mt-tRNA-Gln-TTG-1 (position 9) in control and LP sperm samples (D). Heart_LP_3 sample is excluded from the mt-tRNA-Leu-TAA-1 and mt-tRNA-Thr-TGT-1 analyses due to insufficient coverage (<20 reads).

