## Supplementary figures for "A survey of dietary effects on tRNA abundance and modifications"

#### Supplementary Figure 1

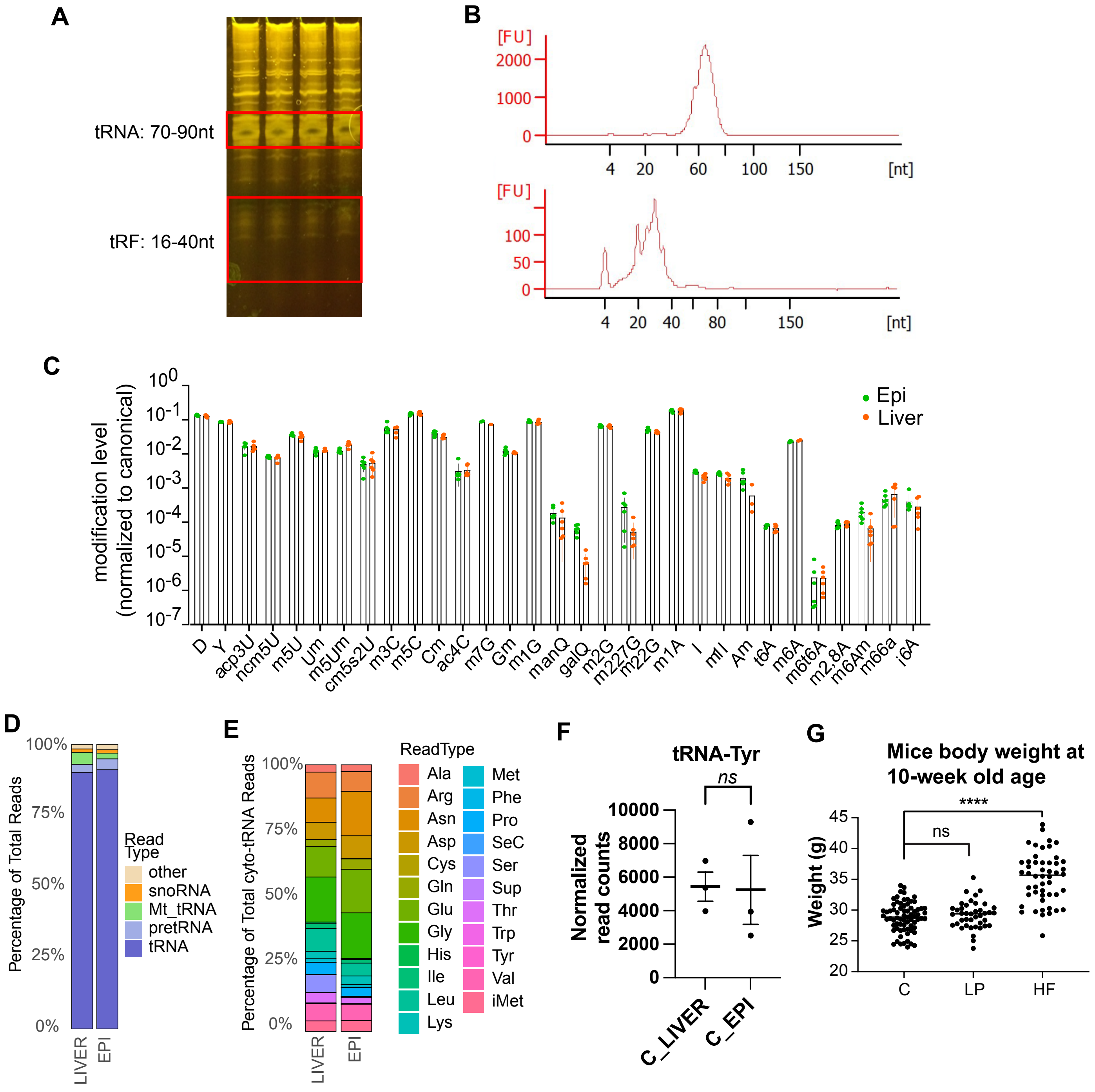

### Supplementary Figure 2

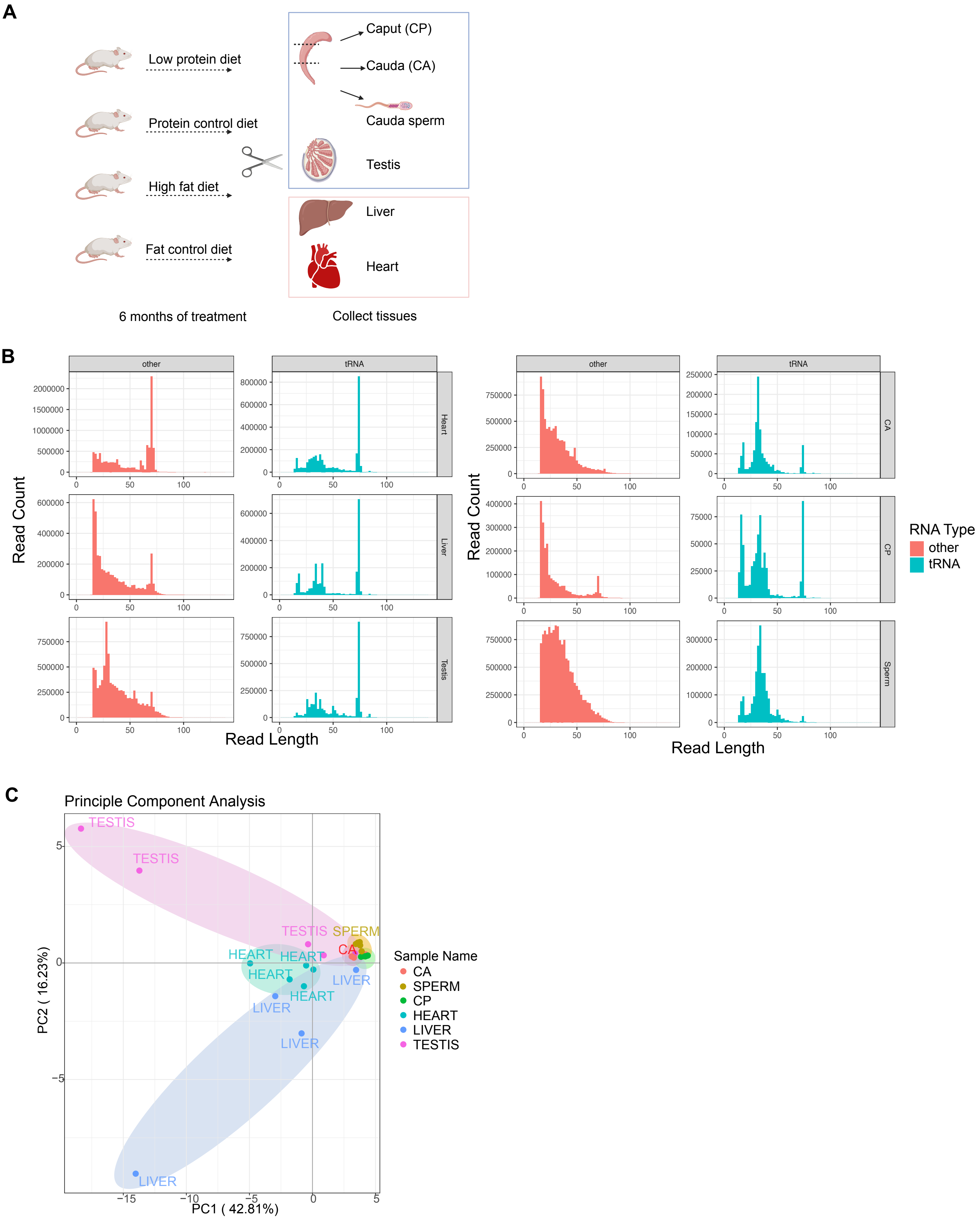

### Supplementary Figure 3

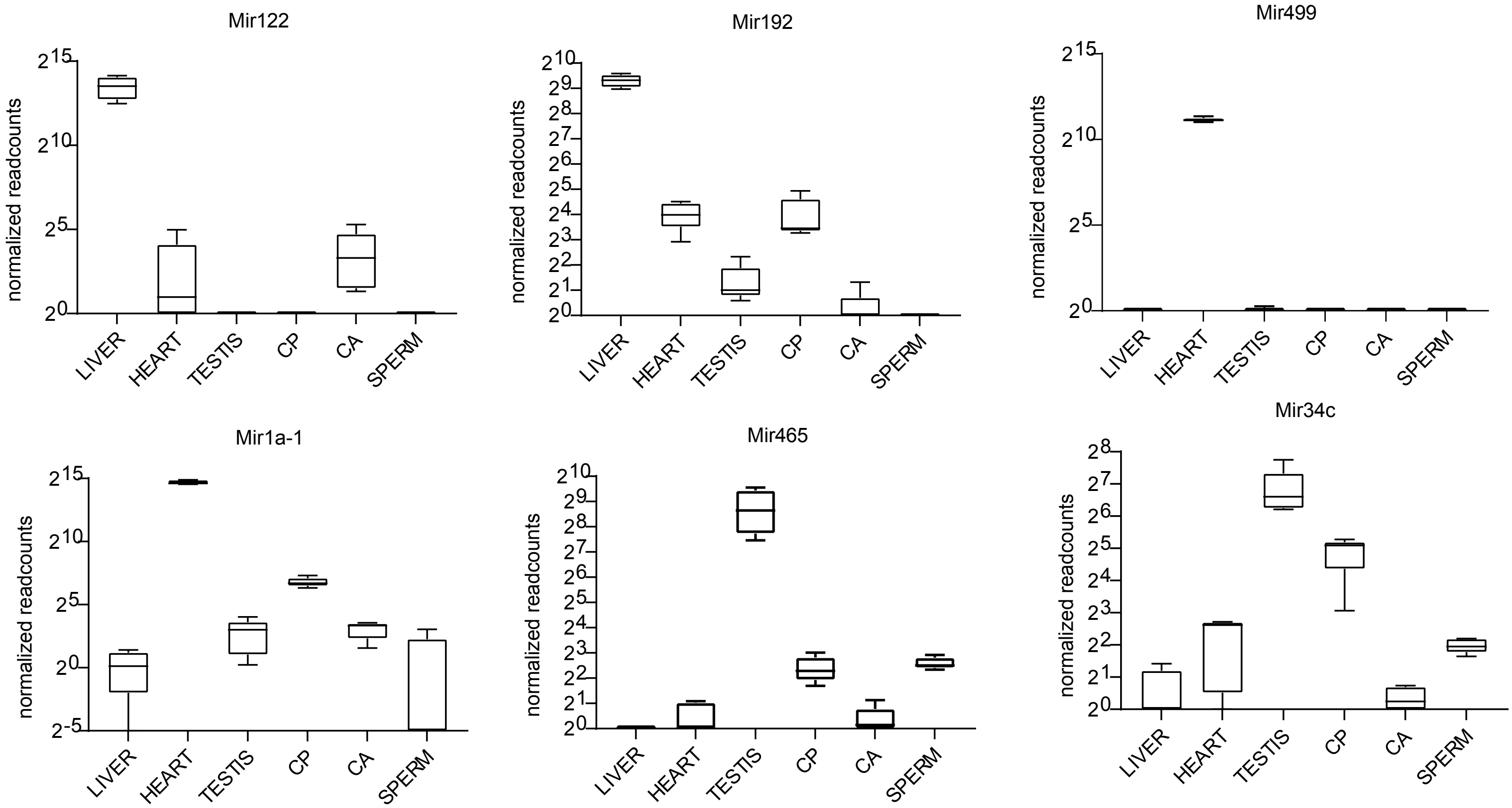

### Supplementary Figure 4

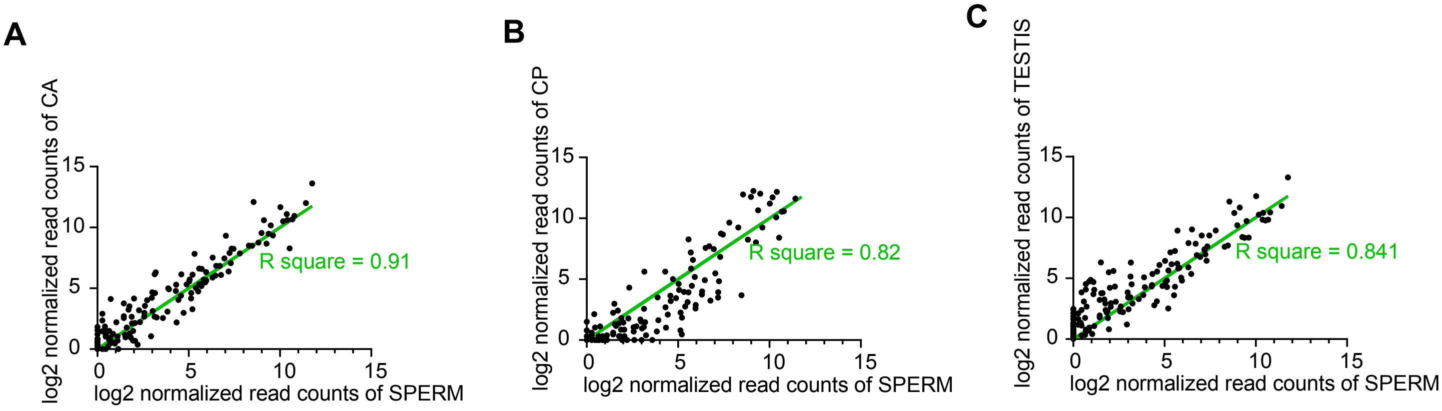

### Supplementary Figure 5

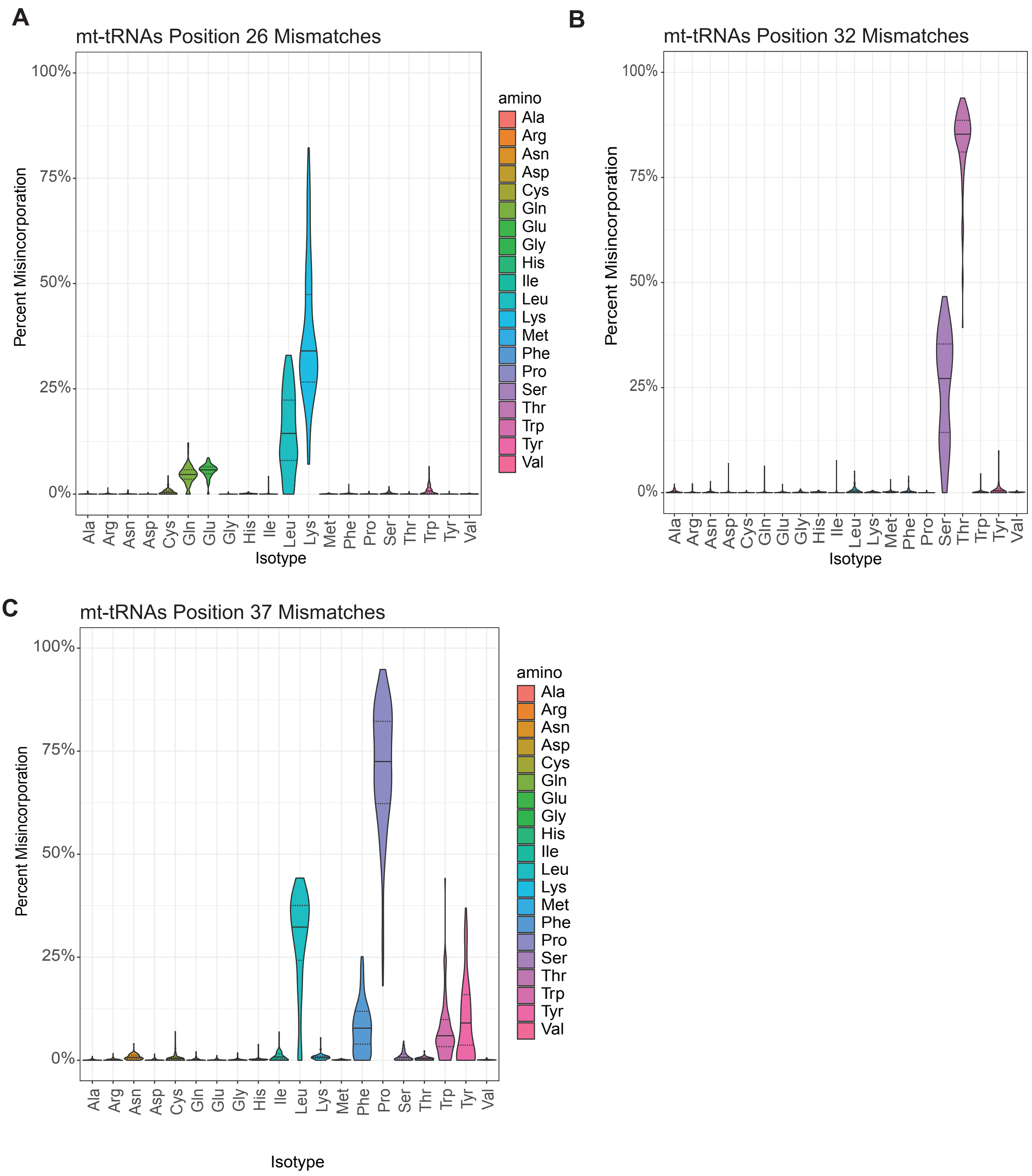

#### Supplementary Figure 6

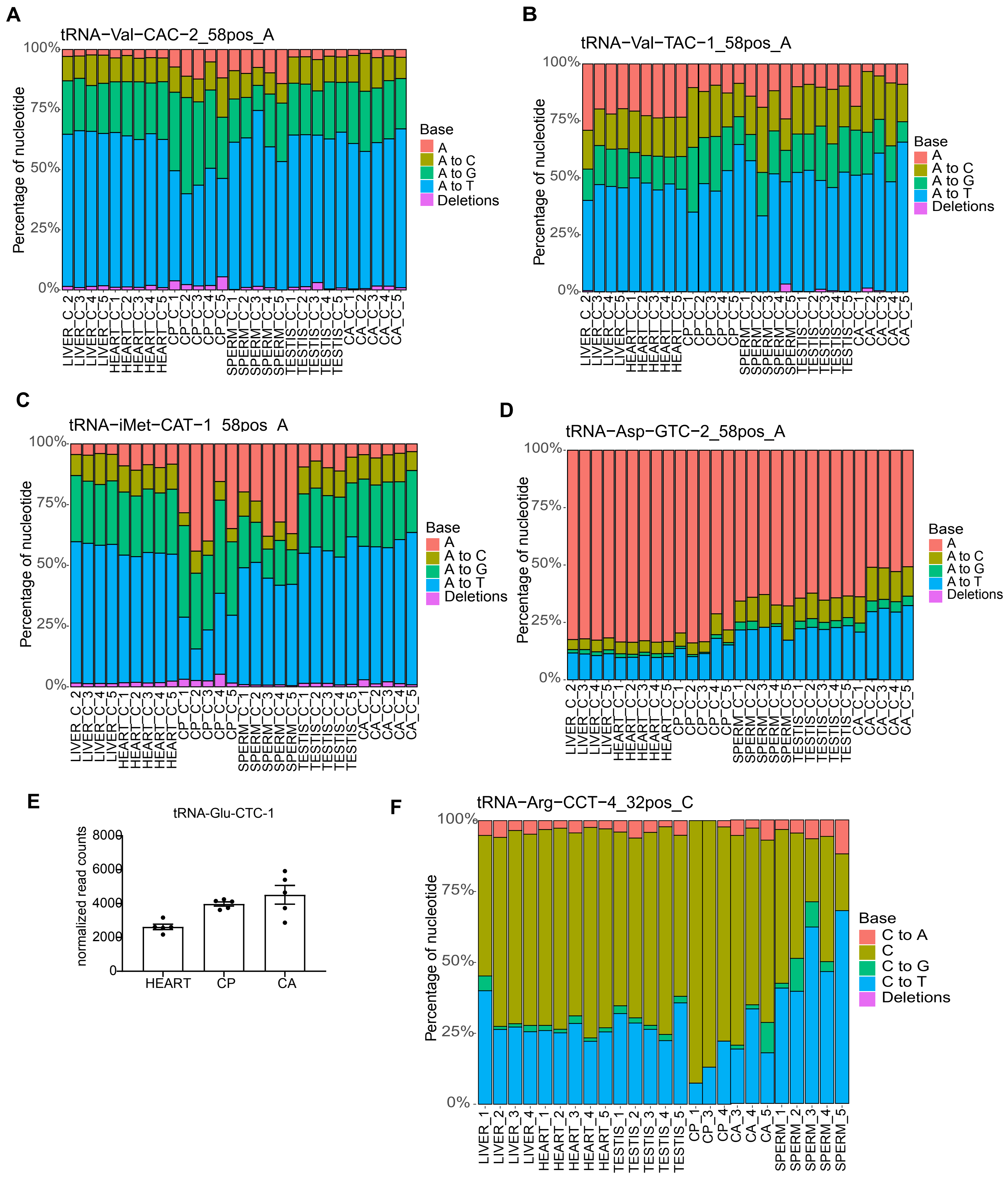

### Supplementary Figure 7

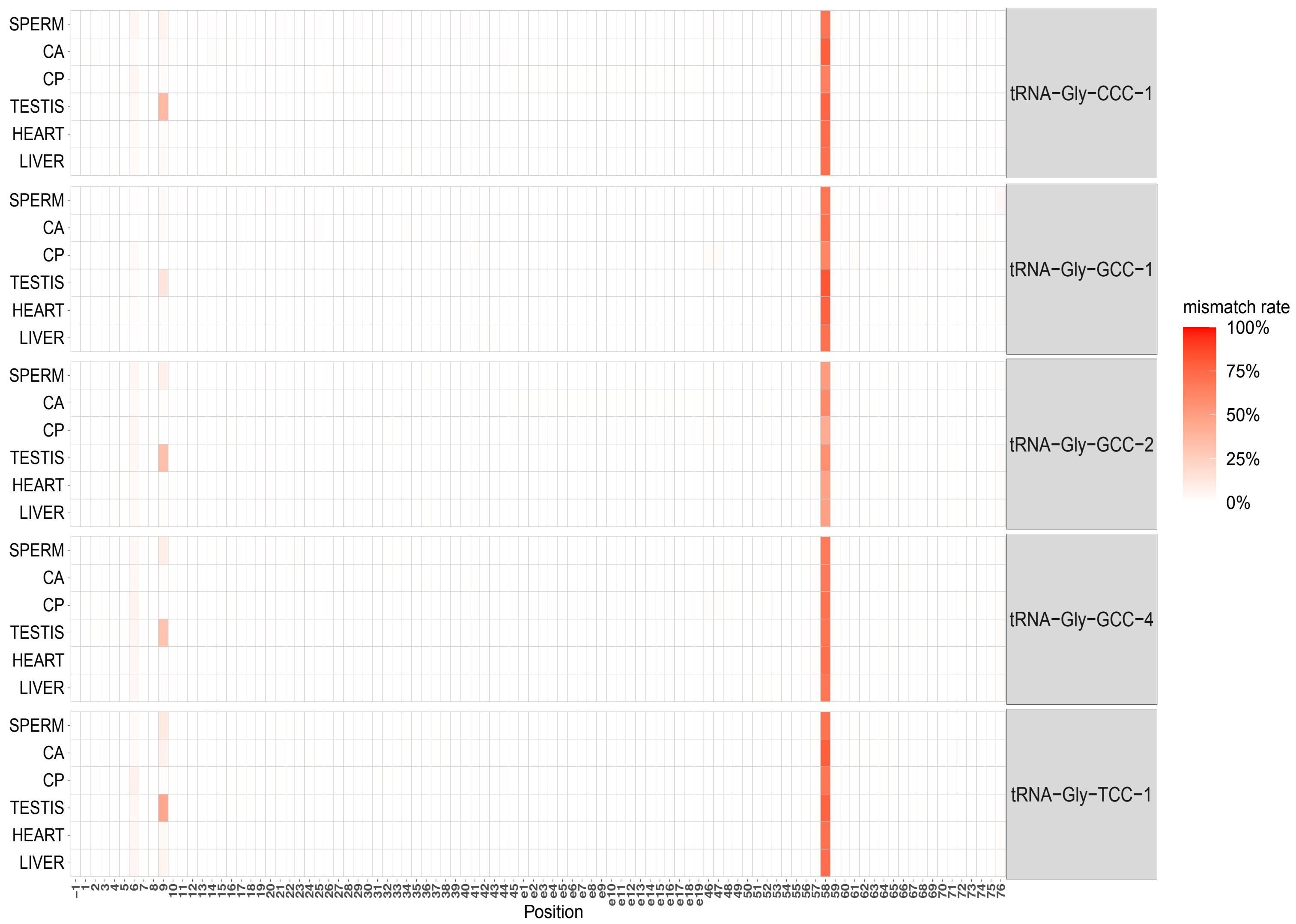

#### Supplementary Figure 8

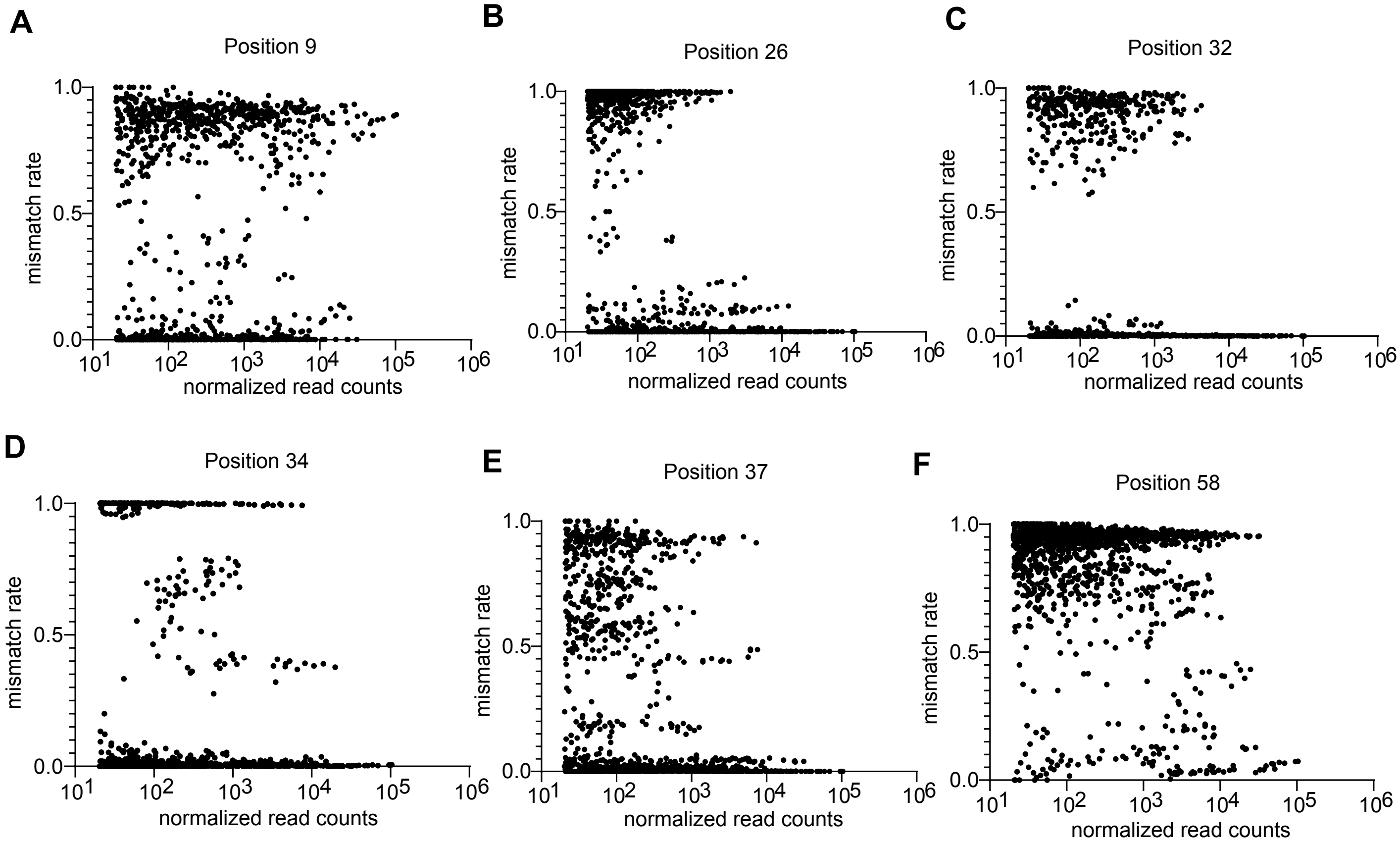

### Supplementary Figure 9

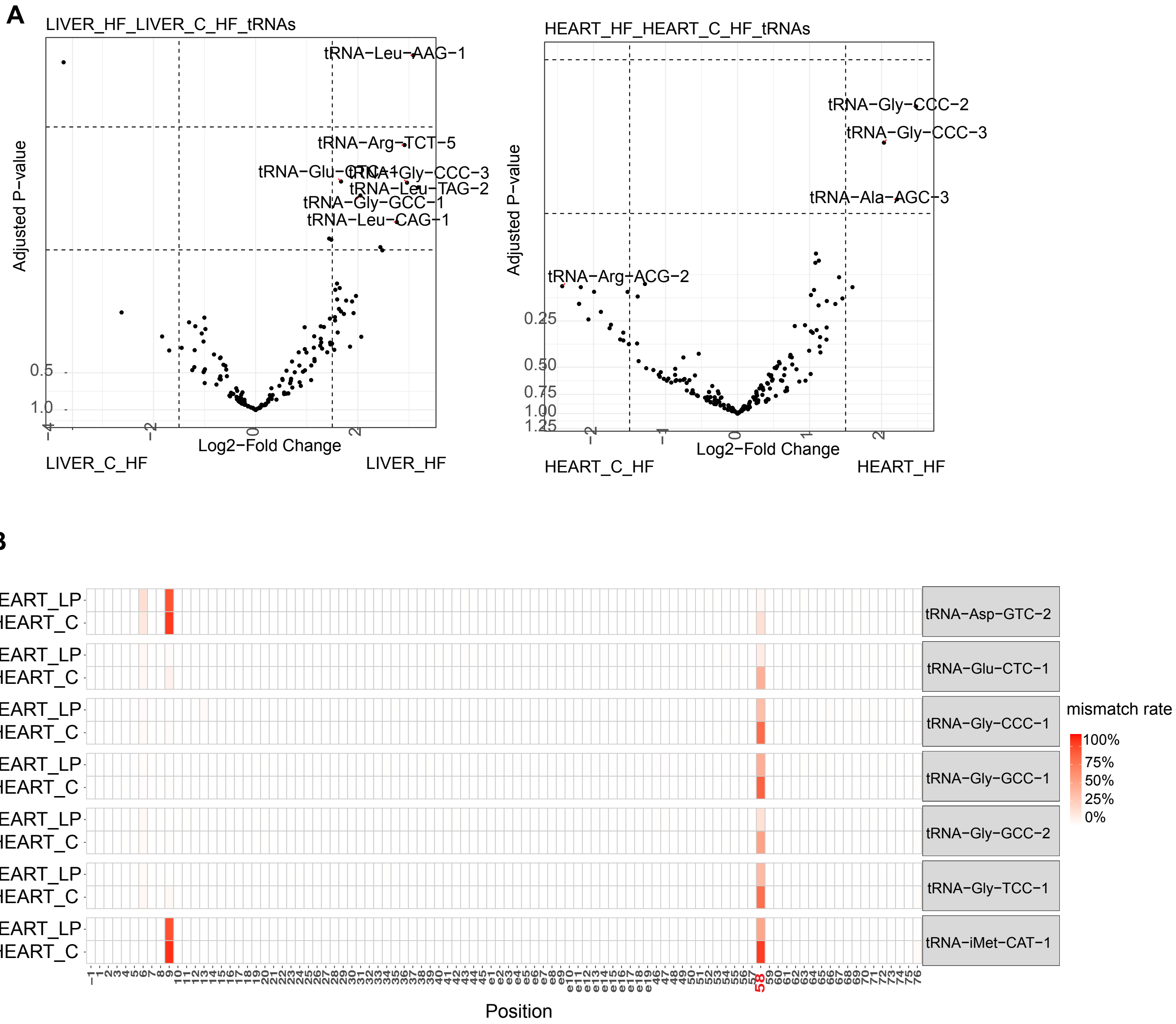

### Supplementary Figure 10

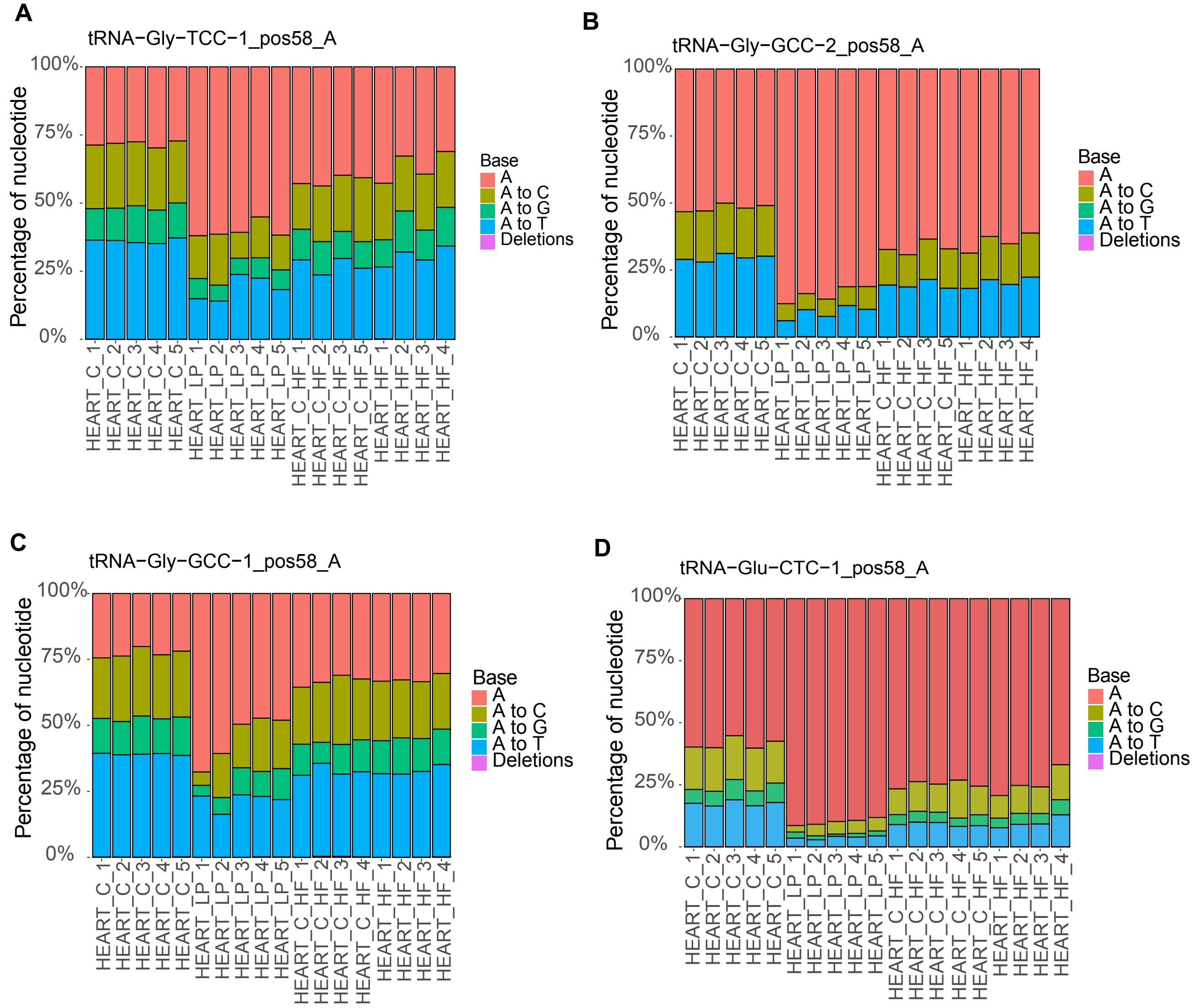

### Supplementary Figure 11

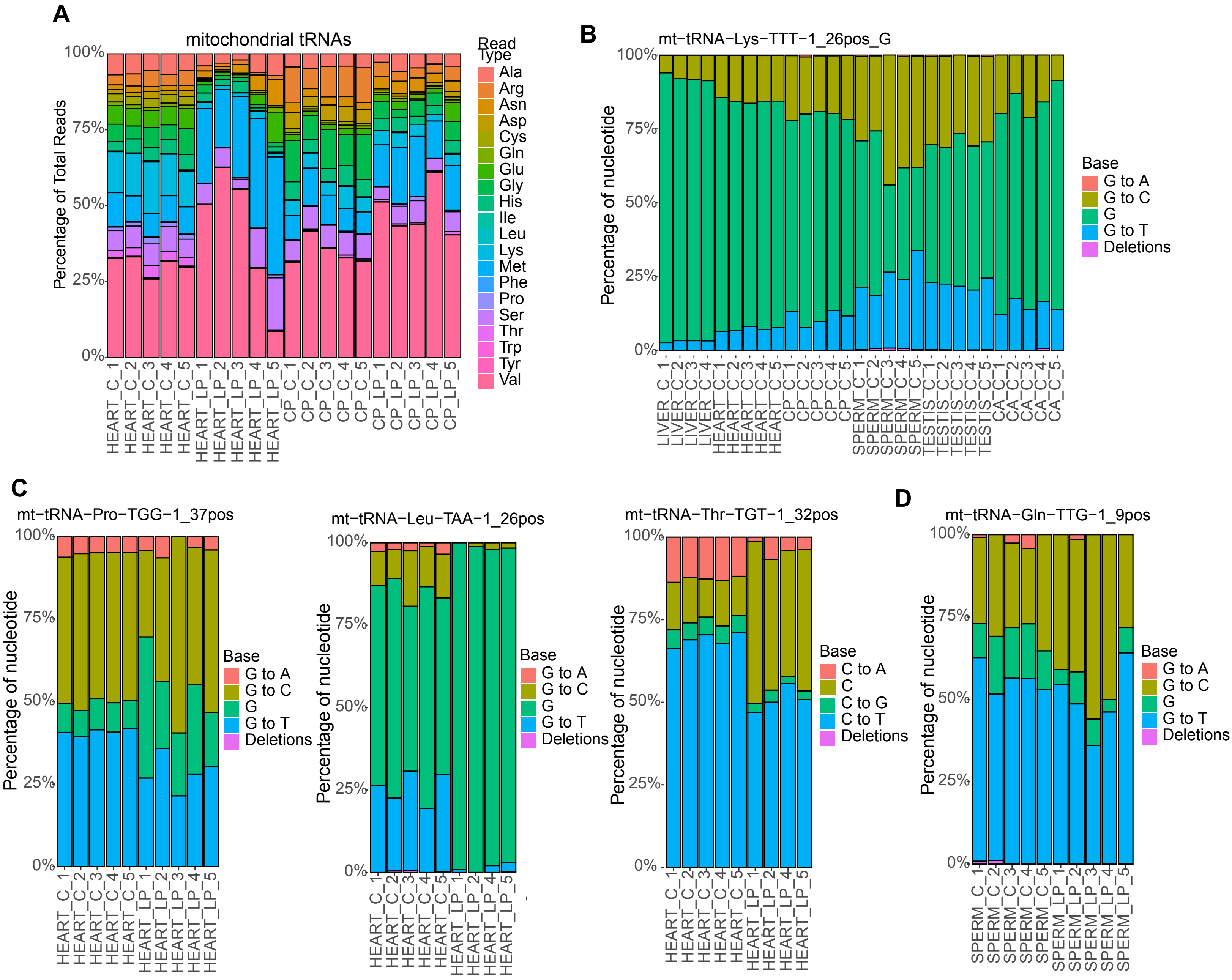
